# Comparing open-source toolboxes for processing and analysis of spike and local field potentials data

**DOI:** 10.1101/600486

**Authors:** Valentina A. Unakafova, Alexander Gail

## Abstract

Analysis of spike and local field potential (LFP) data is an essential part of neuroscientific research. Today there exist many open-source toolboxes for spike and LFP data analysis implementing various functionality. Here we aim to provide a practical guidance for neuroscientists in the choice of an open-source toolbox best satisfying their needs. We overview major open-source toolboxes for spike and LFP data analysis as well as toolboxes with tools for connectivity analysis, dimensionality reduction and generalized linear modeling. We focus on comparing toolboxes functionality, statistical and visualization tools, documentation and support quality. To give a better insight, we compare and illustrate functionality of the toolboxes on open-access dataset or simulated data and make corresponding MATLAB scripts publicly available.

## 1 INTRODUCTION

Analysis of spike and local field potential (LFP) data is an essential part of neuroscientific research (Brown et al., 2004; Stevenson and Kording, 2011; Mahmud and Vassanelli, 2016). There are many already implemented open-source tools and toolboxes for spike and LFP data analysis. However, ascertaining whether functionality of the toolbox fits users’ requirements is in many cases time-consuming. Often neuroscientists are even not aware that some functionality is already implemented and start writing their own scripts from scratch which takes time and is error-prone. We aim to provide a practical guidance for choosing a proper toolbox on the basis of toolbox functionality, statistical and visualization tools, programming language, availability of graphical user interface, support and documentation quality. Compared to the existing reviews (Ince et al., 2009, 2010; Ince, 2012; Mahmud and Vassanelli, 2016; Timme and Lapish, 2018), we

- include in the comparison important toolboxes and tools not covered by earlier reviews (e.g., Brainstorm, Elephant and FieldTrip);
- compare in detail common and discuss unique functionality of toolboxes;
- compare and illustrate functionality of the toolboxes on open-access datasets (Perich et al., 2018; Lawlor et al., 2018; Lowet et al., 2015) and simulated data. For readers’ convenience we make the corresponding MATLAB scripts publicly available^1^;
- overview specialized tools for dimensionality reduction and generalized linear modeling as they are widely used in neuroscientific research (Cunningham and Byron, 2014; Truccolo et al., 2005);
- provide information about documentation and support quality for the toolboxes;
- indicate bibliometric information^2^: while popularity among users alone does not guarantee quality, it can be an important indicator that toolbox’s functions are easy-to-use and have been tested.

### Scope

We include into our comparison major open-source^3^ toolboxes (see Table 1) for spike and LFP data processing and analysis which have a valid link for downloading, documentation, scientific paper describing toolbox’s features or corresponding method, and which were updated during the last five years. In Table 1 we provide a summary of the toolboxes we consider, we list all toolboxes with a brief description in alphabetical order in Section 7 with paper reference and downloading link.

**Table 1:** Features of open-source toolboxes regarding graphical user interface (GUI), visualization tools, Import/Export of spike and LFP data in various file formats, e.g. recorded with different software/hardware, principal programming language, availability of documentation, number of citations, and support by updates at least once per year.

| Toolbox, version | GUI | Visualization | Import/<br>Export | Language | Docu-<br>mentation | Cited | Support |
| --- | --- | --- | --- | --- | --- | --- | --- |
| Brainstorm 3.4 | + | + | + | MATLAB | + | >1000 | + |
| Chronux 2.12 v03 | - | + | + | MATLAB | + | >300 | In part |
| Elephant v0.6.0 | - | - | In part | Python | In part | <30 | + |
| FieldTrip 23.11.18 | - | + | + | MATLAB | + | >3000 | + |
| gramm 2.25 | - | + | - | MATLAB | + | <30 | + |
| Spike Viewer 0.4.2 | + | + | In part | Python | In part | <30 | + |
| SPIKY 3.0 | + | + | - | MATLAB<br>Python | In part | <30 | In part |

Considered toolboxes were developed in MATLAB^4^ and Python^5^ languages which are popular in neuroscientific community.

We have not listed in Table 1 toolboxes FIND (Meier et al., 2008), infotoolbox (Magri et al., 2009) and STAtoolkit (Goldberg et al., 2009), since they are not available under the links provided by the authors (accessed on 27.03.2019); toolboxes BSMART (Cui et al., 2008), DATA-Means (Bonomini et al., 2005), MEA-tools (Egert et al., 2002), MEAbench (Wagenaar et al., 2005), sigTOOL (Lidierth, 2009), SPKTool (Liu et al., 2011), STAR (Pouzat and Chaffiol, 2009), since they have not been updated during the last five years (since 2008, 2005, 2007, 2011, 2011, 2011, and 2012, correspondingly); toolbox SigMate (Mahmud et al., 2012) since it is in beta version; and toolbox OpenElectrophy (Garcia and Fourcaud-Trocmé, 2009) which is not recommended for new users by the toolbox authors^6^.

### Documentation/Support

We have indicated “In part” in Documentation column for Spike Viewer and SPIKY since, compared to other toolboxes from Table 1, they do not provide a description of input parameters for most of the functions. This complicates understanding of implementation details for programming-oriented users that use only a part of the toolbox functionality in their analysis workflow. gramm toolbox specifies function input parameters not in code comments but in separate documentation file^7^. Considered version of Elephant provides only getting started tutorial, more tutorials are to be added^8^. Chronux and SPIKY (MATLAB version) toolboxes are not uploaded to GitHub or other public version control systems, which prevents from tracking version differences and smoothly reporting bugs (Python version of SPIKY is on GitHub^9^).

### Import/Export

We have indicated “In part” in Import/Export column for Elephant and Spike Viewer toolboxes since they require Neo-based Python package^10,11^ (Garcia et al., 2014) for the support of spike file formats (Spike2, NeuroExplorer, AlphaOmega, Blackrock, Plexon etc.). This Neo-based package is popular in neuroscientific society but requires either a separate installation or data conversion to Neo-compatible data format. Brainstorm, Chronux and FieldTrip support working with several spikes file formats (e.g. Blackrock, CED, Neuralynx, Plexon etc.)^12^ as well as working with data from MATLAB workspace or stored as .mat files. SPIKY and gramm support working with data from MATLAB workspace; SPIKY also supports working with data stored in .mat and .txt file formats.

### Compatibility

Chronux under Macintosh operating system requires recompilation of the locfit^13^ and spikesort packages. All other listed toolboxes are supported by Microsoft Windows, Macintosh and Linux operating system. Chronux, FieldTrip, gramm and SPIKY require MATLAB installation, Elephant requires Python installation, Brainstorm and Spike Viewer require neither MATLAB nor Python installation.

### Test dataset

We consider for illustration of toolboxes functionality an open-access dataset (Lawlor et al., 2018; Perich et al., 2018) and refer to this dataset further as “test dataset”. The dataset contains extracellular recordings from premotor (PMd) and primary motor (M1) cortex from a macaque monkey in a sequential reaching task where monkey controlled a computer cursor using arm movements. A visual cue specified the target location for each reach. The monkey receives a reward after making four correct reaches to the targets within the trial.

In Sections 2 and 3, we compare toolboxes for the general spike and LFP data analysis, correspondingly. In Section 4, we compare tools for the analysis of synchronization and connectivity in spike and LFP data. Each of Sections 2-4 is subdivided into two subsections: first, we compare common toolboxes functionality, then we discuss unique toolboxes functionality, i.e. functionality implemented only in one of the toolboxes under comparison. In Section 5, we compare toolboxes with specialized tools for dimensionality reduction and generalized linear modeling. Finally, we summarize the comparisons in Section 6. In Section 7, we list all the considered toolboxes in alphabetical order with links for toolbox downloading and brief descriptions. We do not consider in this review toolboxes specializing on spike sorting and modeling spiking activity. For this we refer to (Ince et al., 2010; Mahmud and Vassanelli, 2016) and web-reviews^14,15,16^ correspondingly.

## 2 TOOLBOXES FOR SPIKE DATA PROCESSING AND ANALYSIS

In Table 2 we compare major open-source toolboxes for spike data analysis, both for point-process data and for spike waveforms. Functionality related to synchronization and connectivity analysis (e.g. cross-correlation, coherence, joint peri-stimulus time histogram, spike-LFP phase-coupling and dissimilarity measures etc.) will be covered in Section 4, and functionality related to dimensionality reduction and generalized linear modeling in Section 5.

**Table 2:** Comparing open-source spike data processing and analysis toolboxes. CV2 – measure of inter-spike variability (Holt et al., 1996), ISIH – Inter-Spike Interval Histogram, locfit – local regression and likelihood based analyses (Bokil et al., 2010; Loader, 2006), LV – measure of Local Variation (Shinomoto et al., 2003), MTF – MultiTaper Fourier transform for point-process data (Jarvis and Mitra, 2001; Bokil et al., 2010), PSTH – Peri-Stimulus Time Histogram

| Toolbox | ISIH | PSTH | Raster plots | Spike sorting | Tuning curves | Statistical tools | Unique features |
| --- | --- | --- | --- | --- | --- | --- | --- |
| Brainstorm | – | – | + | + | + | + | – |
| Chronux | + | + | – | – | – | + | locfit, MTF |
| Elephant | + | + | – | – | – | + | CV2, Fano factor, LV |
| FieldTrip | + | + | + | + | – | + | waveform statistics |
| gramm | – | + | + | – | + | + | – |
| Spike Viewer | + | + | + | – | – | – | – |
| SPIKY | – | + | + | – | – | – | – |

From Table 1 and 2 one can see that Brainstorm, Chronux and FieldTrip toolboxes provide more versatile functionality (see also below) than others, are highly cited, well-documented and allow import from many file formats. The Elephant toolbox has versatile functionality (see Subsection 2.2) but it does not have built-in visualization tools (Elephant provides visualization examples in the documentation using `matlabplot` Python library). Compared to other toolboxes from Table 2,

- Brainstorm and FieldTrip include detailed documentation with tutorials and examples (documentation of other toolboxes from Table 2 has less examples/tutorials for spike data analysis) and have either a forum^17^ or a discussion list^18^ where users can ask questions on data analysis; both toolboxes regularly hold hands-on courses^19,20^, while other toolboxes from Table 2 provide neither forums nor courses;
- Brainstorm and FieldTrip are actively developing by including new functionality;
- FieldTrip provides many descriptive and inferential statistics mostly not requiring MATLAB statistical toolbox (Brainstorm provides statistical tools^21^ without examples for spike data analysis^22^ and these statistical functions are not part of spike data analysis functions, different to how it is often done in FieldTrip and Chronux; Spike Viewer and SPIKY do not provide statistical tools for general spike data analysis);
- FieldTrip and gramm allow versatile data plots customization (color maps, line widths, smoothing, errorbars etc.); while gramm provides better and quicker general visualization tools, FieldTrip provides plotting customization specific for spike data analysis (conditions/interval/trials/channels and optimal bin size selection);
- for programming-oriented users, Chronux and FieldTrip provide, to our opinion, most convenient and well-commented data analysis pipeline with clear uniform data structure (other toolboxes from Table 2 are lacking at least one of three following components: detailed code comments with description of input/output parameters, uniform data structure throughout the analysis pipeline, modular function design allowing to easily adapt them into analysis workflow). Chronux reference documentation in the function description provides a list of functions which are called from the function and from which the function is called, this is convenient for programming-oriented users.

### 2.1 Comparing common tools: peri-stimulus time-histogram, raster plot, inter-spike interval histogram and spike sorting

In this subsection we compare most common spike data analysis functions: peri-stimulus time histogram (PSTH), raster plot, inter-spike interval histogram (ISIH) and spike sorting algorithms for toolboxes from Table 2. Regarding visualization, the gramm visualization toolbox stands out with its publication-quality graphics, which helps avoiding major post-processing. This is illustrated in Figure 1, where we compare PSTH and raster plots for test dataset produced in FieldTrip and gramm toolboxes, both of which provide most adjustable plot properties compared to other toolboxes from Table 2 (see below a detailed comparison).

**Figure 1:**
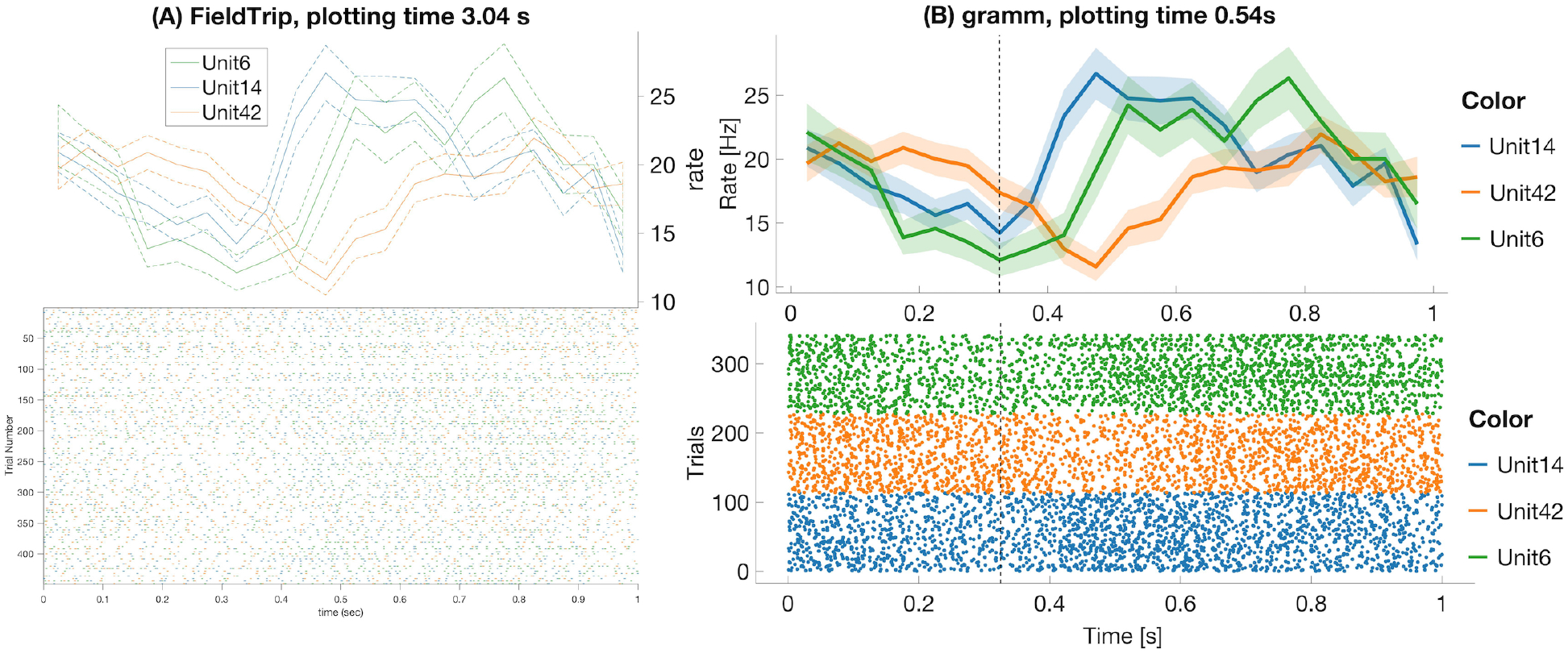
FieldTrip (A) and gramm (B) provide most adjustable peri-stimulus (PSTH) and raster plots properties (plotting time is averaged over 1000 runs, MATLAB 2016a, here and later for processor 3.2 GHz Intel Core i5 with 16GB RAM) among toolboxes from Table 2. We considered 50 ms bin size, M1 units 6, 14, 42, monkey MM for the test dataset. PSTHs are presented with standard error of the mean, neural activity is aligned to trial start for reaches toward the second target in the trial. FieldTrip build-in tools do not allow to adjust font size in a raster plot and line width in a PSTH plot (one has to do it manually with MATLAB tools), and do not allow to plot raster and PSTH in separate figures (though one can plot spike densities in a separate figure). Advantages of gramm toolbox for PSTH and raster plots are quick plotting, raster plots separation for different units, vertical dashed lines for showing event times of the experiment protocol, and smooth adjustment of line width, font size, color maps, errorbar, components positions, etc.

We do not provide raster plots and PSTH plots for other toolboxes from Table 2 with visualization tools since

- Brainstorm does not provide PSTH plots; raster plots are available only for one unit per figure^23^;
- Chronux does not provide raster plots and allows to plot only smoothed PSTH for one unit per figure without built-in tools to adjust line width, font size, colors etc.;
- in SPIKY raster and PSTH plots are available only for one unit per figure without built-in tools to adjust line width, marker size, font style and size, colors (Kreuz et al., 2015, Figure 2) and without confidence intervals for PSTHs;
- in Spike Viewer PSTH plots are available without confidence intervals^24^.

**Figure 2:**
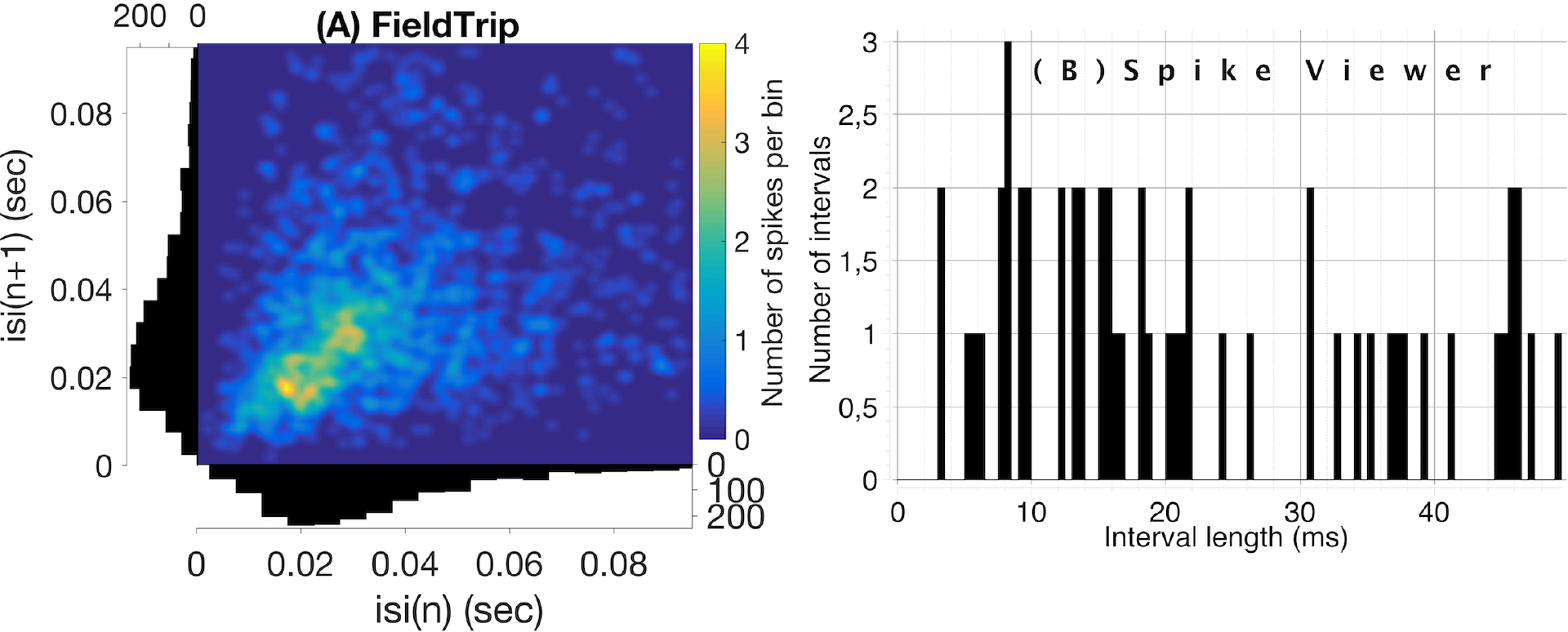
Compared to Spike Viewer (B), FieldTrip (A) provides also a second-order statistic on inter-spike interval histogram (ISIH). We considered test dataset (M1 unit 14 aligned to trial start for reaches towards the first target, monkey MM) for FieldTrip plot and Spike Viewer test dataset for Spike Viewer plot. Font sizes in FieldTrip have been adjusted with MATLAB tools since FieldTrip built-in tools do not provide this option.

Regarding statistical tools when computing PSTHs, Chronux computes PSTH for adaptive or user-defined kernel width with Poisson error or bootstrapped over trials (both with doubled standard deviation error). Elephant computes PSTH for fixed user-defined bin size without additional statistics (note that Elephant provides many kernel functions for convolutions such as rectangular, triangular, Guassian, Laplacian, exponential, alpha function etc.). FieldTrip computes PSTH for optimal (by Scott’s formula (Scott, 1979)) or user-defined bin width with variance computed across trials. Besides, FieldTrip, different to other toolboxes from Table 2, allows statistical testing on PSTHs for different conditions or subjects^25^ with a parametric statistical or a non-parametric permutation test. Brainstorm provides this functionality by calling FieldTrip functions. gramm allows to compute PSTHs with (bootstrapped) confidence intervals, standard error of the mean, standard deviation etc^26^ only for user-defined bin width. Spike Viewer and SPIKY compute PSTH only for user-defined bin width and do not compute statistics for PSTHs across trials.

In Figure 2 we compare visualization of ISIH provided by FieldTrip and Spike Viewer since other toolboxes from Table 2 do not provide ISIH visualization (Brainstorm, Chronux and Elephant compute ISIH without visualization, see details below).

Regarding statistical tools when computing ISIH, FieldTrip computes ISIH with a coefficient of variation (a ratio of the standard deviation to the mean), Shinomoto’s local variation measure (Shinomoto et al., 2005) or a shape scale for a gamma distribution fit. Chronux computes ISIH with two standard deviations away from the mean calculated using jackknife resampling. Elephant computes ISIH with a coefficient of variation. Spike Viewer does not compute statistics on ISIH.

Brainstorm and FieldTrip provide spike sorting algorithms including spike detection and extraction, i.e., using time-continuous broadband data as input. Spike sorting package is no longer provided by Chronux. Brainstorm implements supervised and unsupervised spike sorting according to the methods WaveClus (Quiroga et al., 2004), UltramegaSort2000 (Hill et al., 2011; Fee et al., 1996), KiloSort (Pachitariu et al., 2016) and Klusters (Hazan et al., 2006). FieldTrip implements k-means and Ward (for several Ward distances) sorting methods. While Chronux and FieldTrip do not provide tutorials on spike sorting, Brainstorm has a detailed tutorial^27^.

Brainstorm provides computing and visualization of tuning curves: they are plotted with one figure per unit for selected units, conditions and time interval but without customization of font size, line width and colors, no variance statistic across trials is computed^28^. gramm toolbox provides visualization of tuning curves including fits from MATLAB curve smoothing toolbox and user-defined functions (also in polar coordinates) with (bootstrapped) confidence intervals, standard error of the mean, standard deviation etc. As the considered gramm version is not focused on spike data analysis, firing rates averaged per condition need to be computed prior to tuning curves visualization (see example in our open MATLAB script).

### 2.2 Description of unique tools

In this subsection we discuss unique tools of toolboxes from Table 2, e.g. fitting tools, and higher order statistics (variability and spectral measures) on spike timing.

Chronux provides two unique tools: local regression package (locfit) and point-process spectrograms. locfit is based on local regression methods (Loader, 2006; Parikh, 2009; Hayden et al., 2009) and provides a set of methods for fitting functions and probability distributions to noisy data. The idea of local regression is that the estimated function is approximated by a low order polynomial in a local neighborhood of any point with polynomial coefficients estimated by the least mean squares method (Bokil et al., 2010). In (Bokil et al., 2010; Loader, 2006) local regression methods are motivated by their simplicity, non-parametric approach to kernel smoothing and by reducing the bias at the boundaries which is present in kernel smoothing methods. On the other hand, it was shown that fixed and variable kernel methods (Shimazaki and Shinomoto, 2010, Algorithm 2, Appendix A.2) as well as Abramson’s adaptive kernel method (Abramson, 1982) outperform `locfit` for simulated data examples (Shimazaki and Shinomoto, 2010).

Point-process spectrograms are usually used to illustrate rhythmic properties of otherwise stochastic spiking patterns rather than for statistical inference (Deng et al., 2013). We refer to (Hurtado et al., 2004, 2005) regarding methods to evaluate statistical significance of point-process spectral estimators and to (Jarvis and Mitra, 2001; Rivlin-Etzion et al., 2006) for a critical discussion. Chronux provides the only open-source, to our knowledge, implementation of point-process spectral estimates which is implemented according to (Jarvis and Mitra, 2001; Rivlin-Etzion et al., 2006, Section 4, Formula 11), see example of usage in our open MATLAB script.

Elephant provides several statistical measures for spike timing variability such as Fano factor, CV2 measure of inter-spike variability (Holt et al., 1996) and a measure of local variation (Shinomoto et al., 2003) which were introduced as substitutes of classical coefficient of variation to overcome its sensitivity to firing rate fluctuations between trials (Shinomoto et al., 2005).

FieldTrip allows to compute mean average spike waveform and its variance across trials, one can optionally align waveforms based on their peaks, rejects outlier waveforms and interpolate the waveforms.

## 3 TOOLBOXES FOR LFP DATA ANALYSIS

In Table 3 we compare open-source toolboxes for processing and analysis of local field potential (LFP) data. Functionality related to synchronization and connectivity analysis will be discussed in Section 4.

**Table 3:**
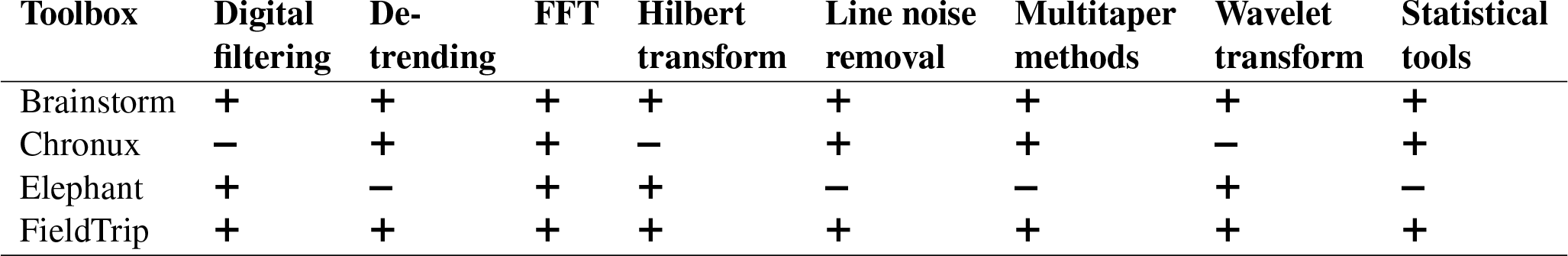
Comparing open-source toolboxes for processing and analysis of LFP data. FFT – Fast Fourier Transform

From Table 3 one can see that Brainstorm and FieldTrip toolboxes provide most versatile functionality for LFP data analysis. Compared to other toolboxes from Table 3,

- FieldTrip provides most flexible and versatile digital filtering (in particular, a fast and accurate line noise removal technique) and spectral analysis tools (see details in Subsection 3.1);
- Brainstorm^29,30^ and FieldTrip^31,32^ provide detailed tutorials with guidance on parameter choice and examples for digital filtering and spectral analysis. Chronux provides examples on parameter choice for spectral analysis in manuals^33^ (Pesaran, 2008);
- Brainstorm and Elephant provide fast implementation of Morlet wavelet transform (see details in Subsection 3.1);
- Brainstorm, Chronux and FieldTrip provide statistical tools for computing variance across trials and for comparing between conditions when estimating spectra; Elephant does not compute statistics on the estimated spectra;
- Brainstorm and FieldTrip allow adjustment of plot properties for spectral analysis such as baseline correction, trials and channels selection, colormaps and interactive selection of spectrogram part for further processing. Neither Chronux nor Elephant provide these options. Compared to Brainstorm, FieldTrip also allows to adjust font sizes, titles, plot limits etc.

### 3.1 Comparing common tools: filtering, detrending and spectral analysis

Digital filtering is implemented in Brainstorm, FieldTrip and Elephant toolboxes. Compared to toolboxes from Table 3, MATLAB and Python themselves provide more flexible filtering tools. Yet, it is convenient to have filtering within the toolbox pipeline. First, it allows to avoid extra conversion from toolbox’s format to MATLAB/Python and back. Second, toolboxes allow simplified setting of filter parameters for typical neuroscientific datasets and offer tutorials for their choice for non-experienced users.

Brainstorm, FieldTrip and Elephant toolboxes provide low/high/band-pass and band-stop filters for user-defined frequencies.

- Brainstorm provides Finite Impulse Response (FIR) filters with Kaiser window based on kaiserord functions from MATLAB Signal Processing Toolbox (Octave-based alternatives are used if this toolbox is not available). The user can set 40 or 60 dB stopband attenuation, data are padded with zeros at edges with a half of filter order length (according to the description of the filtering `bst_bandpass_hfilter` function used by default);
- Elephant provides Infinite Impulse Response (IIR) Butterworth filtering with adjustable order using `scipy.signal.filtfilt` (with default padding parameters) or `scipy.signal.lfilter` standard Python functions;
- FieldTrip provides the most flexible filtering tools with user-defined filter type (Butterworth IIR, window sinc FIR filter, FIR filter using either standard MATLAB `fir1` or `firls` function from Signal Processing Toolbox or frequency-domain filter using standard `fft` and `ifft` MATLAB functions), padding type and optional parameters such as window type (Hanning, Hamming, Blackman, Kaiser), filter order and direction, transition width, passband deviation, stopband attenuation etc.^34^. An automatic tool to deal with filter instabilities (which MATLAB 2016a, to our knowledge, does not provide) is implemented by either recursively reducing filter order or recursively splitting the filter into sequential filters.

Brainstorm, Chronux and FieldTrip also provide specific tools for line noise removal. Brainstorm reduces line noise with IIR notch filter (employing either `filtfilt` function from MATLAB Signal Processing toolbox or MATLAB `filter` function). Chronux reduces line noise using Thomson’s regression method for detecting sinusoids (Thomson, 1982). FieldTrip reduces line noise by two alternative methods: with a discrete Fourier transform (DFT) filter (by fitting a sine and cosine at user-defined line noise frequency and subsequently subtracting estimated components) or by spectrum interpolation (Mewett et al., 2004). In Figure 3 we compare 60 Hz line noise removal by Chronux, FieldTrip and Brainstorm toolboxes on the basis of an example provided by MATLAB^35^ for open-loop voltage across the input of an analog instrument in the presence of 60 Hz power-line noise. One can see that FieldTrip selectively and successfully attenuates 60 Hz while Brainstorm does not fully suppress 60 Hz, Chronux suppresses also frequencies around 62 Hz, the MATLAB solution contains some remaining oscillations in the beginning of the signal, which is also reflected in the periodogram by a slight inaccuracy around 61-62 Hz. In Figure 3 (C) we present mean squared error (MSE) between power spectrum values of the original and estimated signal except the values estimated in 0.2 Hz vicinity of 60 Hz.

**Figure 3:**
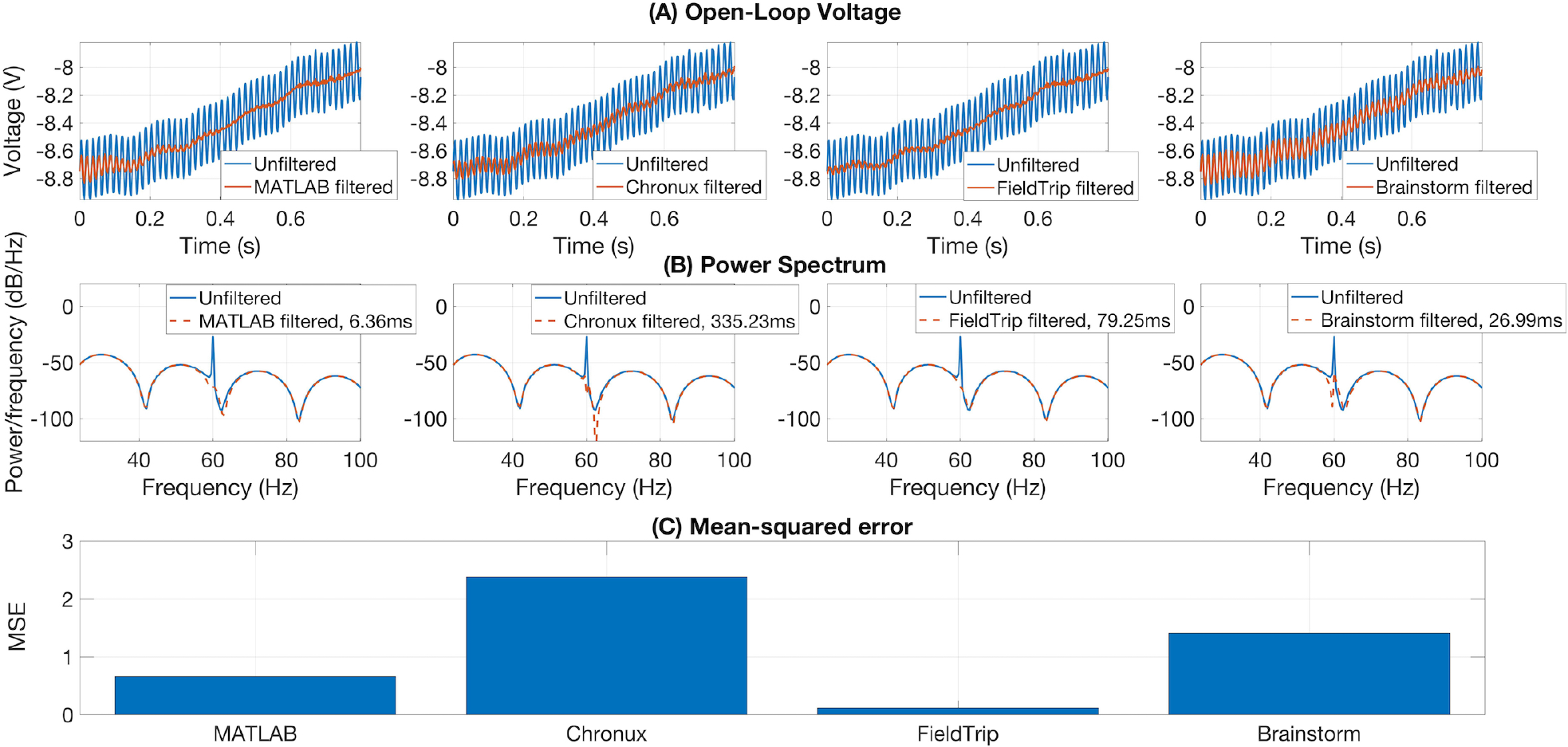
FieldTrip (discrete Fourier transform filter, default parameters) provides the fastest and the most accurate line noise removal compared to MATLAB solution (Butterworth notch filter with 2 Hz width), Chronux (default 5 tapers, bandwidth 3) and Brainstorm (IIR notch filter with 1 Hz width). Filtering times are averaged over 1000 runs, MATLAB 2016a.

Brainstorm, Chronux and FieldTrip provide detrending tools. Brainstorm removes a linear trend from the data, Chronux detrending employs local linear regression^36^, whereas FieldTrip detrending uses a general linear model approach and removes mean and linear trend from the data (by fitting and removing an *N*th order polynomial from the data)^37^: Brainstorm, Chronux and FieldTrip offer similar performance in terms of processing time and trend removal accuracy for a simple MATLAB example^38^ (see our open MATLAB code).

Compared to the classic Fourier transform, multitaper methods provide more convenient control of time and frequency smoothing (Percival and Walden, 1993; Mitra, 2007). Spectral decomposition with Morlet wavelets provides a convenient way of achieving a time-frequency resolution trade-off (van Vugt et al., 2007), since it is inherent to the method that wavelets are scaled in time to vary resolution in time and frequency, see (van Vugt et al., 2007) for a comparison of multitaper and wavelet methods and (Bruns, 2004) for a comparison of wavelet, Hilbert and Fourier transform. Equivalent time-frequency trade-offs can also be implemented with short-time Fourier or Hilbert methods via variable-width tapers (Bruns, 2004).

Chronux and FieldTrip provide multitaper power spectrum estimation using Thomson’s method (Thomson, 1982; Percival and Walden, 1993; Mitra and Pesaran, 1999) with Slepian sequences (Slepian and Pollak, 1961). Additionally to this, FieldTrip allows also more conventional tapers (e.g. Hamming, Hanning). In FieldTrip, the user defines frequencies and time interval of interest, width of sliding window and of frequency smoothing. In Chronux, the user defines bandwidth product and number of tapers to be used (see (Prieto et al., 2007) for a discussion of multitapers parameter choice).

Brainstorm, Elephant and FieldTrip implement complex-valued Morlet transform. FieldTrip provides time-frequency transformation using Morlet waveforms either with convolution in the time domain or with the multiplication in the frequency domain. Brainstorm and Elephant implement convolution in the time domain. FieldTrip implements Morlet wavelet transformation methods based on (Tallon-Baudry et al., 1997), the user defines the wavelet width in number of cycles and optionally wavelet length in standard deviations of the implicit Gaussian kernel. In Brainstorm the user sets the central frequency and temporal resolution. Elephant implements Morlet wavelets according to (Le Van Quyen et al., 2001; Farge, 1992), where the user sets central Morlet frequencies, size of the mother wavelet and padding type.

Different to other toolboxes from Table 3 FieldTrip also implements Fourier transform on the coefficients of the multivariate autoregressive model estimated with FieldTrip tools (see Subsection 4.1 for more details on MVAR implementation in FieldTrip).

Elephant does not compute statistics on estimated power spectrum whereas Chronux and FieldTrip compute confidence intervals and standard error, correspondingly, in a standard way or with jackknife resampling. To compare spectrum estimates for different conditions or subjects, Chronux provides a two-group test and FieldTrip performs a parametric statistical test, a non-parametric permutation test or a cluster-based permutation test (Brainstorm includes these FieldTrip statistical functions).

MATLAB R2016a, compared to Chronux, FieldTrip and Brainstorm,

- does not provide detailed tutorials for multitaper and wavelet parameters choice;
- does not have built-in tools for computing average spectrogram across trials;
- does not have built-in tools for generating multitaper spectrograms;
- uses exclusively short-time Fourier transform for standard spectrogram plotting.

In Figure 4 we compare spectrum estimation methods implemented in Brainstorm (A), Chronux (B), Elephant (C), FieldTrip (D-F) and MATLAB (G-H) for two simulated signals, *x*_1_(*t*) and *x*_2_(*t*).

**Figure 4:**
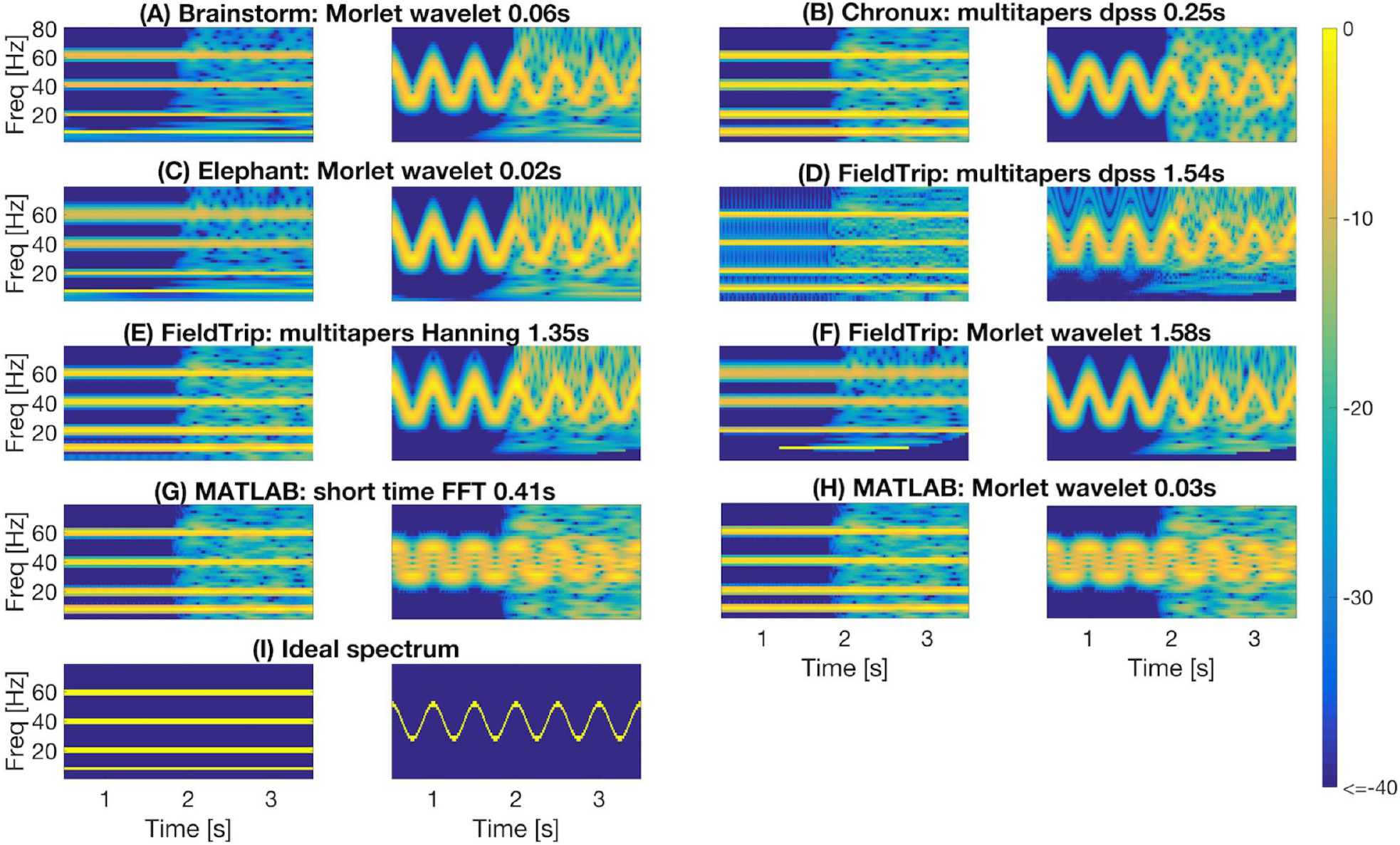
Comparing spectral analysis tools provided by the toolboxes. For each toolbox we plot estimated spectrum of signal *x*_1_ (left subpanel) and of signal *x*_2_ (right subpanel). For short-time FFT we used 0.512 s moving window with 0.001s step. For multitaper methods we used in Chronux a single taper with time bandwidth product 2 (left) and 8 (right); in FieldTrip a single taper with 2 Hz (left) and 0.1*F* (right) frequency smoothing for time window 0.512s (left) and 8*/F* (right) at frequency *F*. For wavelet methods we used in MATLAB and Brainstorm central frequency 4 (left) and 1.5 (right) Hz; in Elephant and FieldTrip 20 (left) and 10 (right) cycles wavelets resulting in the spectral bandwidth *F/*10 (left) and *F/*5 (right) Hz at frequency *F*. Spectrum estimating times were averaged over 1000 runs in MATLAB 2016a.

We generate *x*_1_(*t*) as a sum of sines and *x*_2_(*t*) by sinusoidal frequency modulation, see Eq. 1-2. We add normally distributed pseudo-random values with zero mean to the second half of both signals:

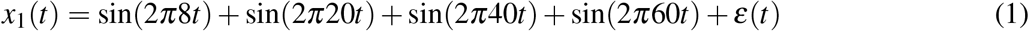

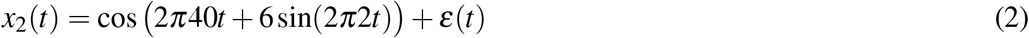

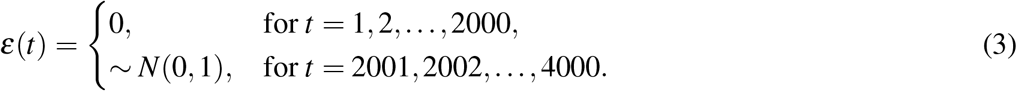

The instantaneous frequency of the signal *x*_2_(*t*) is defined by the following equation (Granlund, 1949):

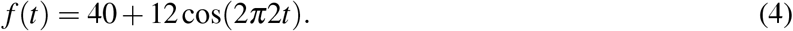

To compare quantitatively the spectra estimated by the toolboxes we compute power spectrum values of the ideal signal by setting maximum spectrum values at theoretical frequencies of the signals *x*_1_ (8, 20, 40 and 60 Hz) and *x*_2_ (given by Eq. 4) and minimum at all other frequencies. When setting ideal power spectrum values we allow bandwidth of 1 Hz, i.e. we set the maximum power spectrum values also at neighboring frequencies. Then we compare in Figure 5 the estimated spectrum values with the ideal spectrum values using mean squared error and two-dimensional Pearson correlation coefficient as suggested in (Rankine et al., 2005).

**Figure 5:**
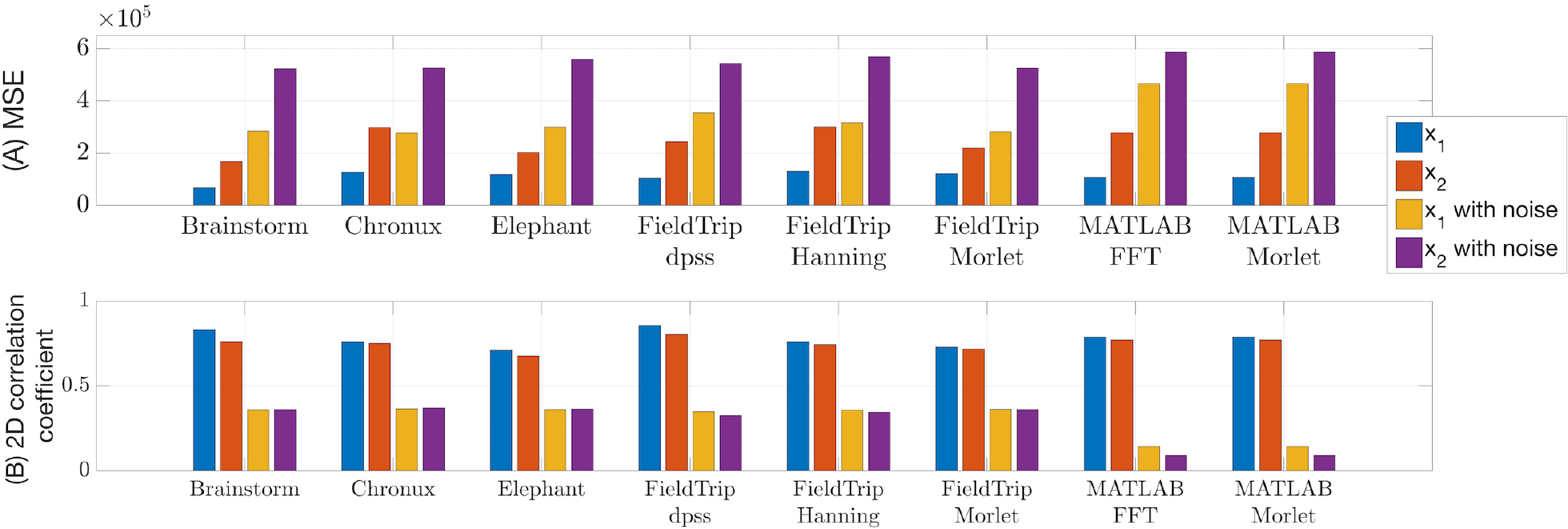
Mean squared error (A) and two-dimensional Pearson correlation coefficient (B) values between estimated and ideal spectra. These measures were computed for the time span from 1 to 3 s for the signals generated according to Eqs. 1-2. The lower MSE and the higher correlation coefficient are, the closer is the estimated spectrum to the ideal spectrum.

From Figures 4-5 we conclude that

- MATLAB standard spectrogram tools are less robust with respect to noise than spectrum estimation provided by the toolboxes from Table 3 for the signal *x*_2_ with changing frequencies;
- while Brainstorm, Chronux, Elephant and FieldTrip provide equally good accuracy of spectra estimation, Brainstorm and Elephant provide the fastest computing tools (see spectra computing times in subplot titles of Figure 4).

See in our open MATLAB script an example of spectral analysis with averaging over trials for real-world LFP data (Lowet et al., 2015).

### 3.2 Description of unique tools

Compared to other toolboxes from Table 3, Chronux provides several unique features for specialized computations (Bokil et al., 2010) such as space-frequency singular value decomposition (SVD) for univariate and multivariate continuous signals: for theoretical details we refer to (Mitra and Pesaran, 1999) and for an example of possible application to (Makino et al., 2017; Prechtl et al., 1997). Space-frequency SVD can be applied to the space-time data as, for example, in (Prechtl et al., 1997), where space-frequency SVD has been applied for spectral analysis of transmembrane potentials optically recorded in pixels distributed in space. Chronux also provides computation of multitaper spectral derivatives and stationarity statistical test for continuous processes based on quadratic inverse theory.

Elephant provides computing of the current source density from LFP data using electrodes with 2D or 3D geometries.

## 4 TOOLBOXES WITH SYNCHRONIZATION AND CONNECTIVITY ANALYSIS TOOLS

In Table 4 we compare open-source toolboxes providing tools for spike-spike, field-field (LFP-LFP) or spike-field (spike-LFP) synchronization and connectivity analysis. We refer to (Blinowska, 2011; Bastos and Schoffelen, 2016) for reviews of functional connectivity analysis methods and their interpretational pitfalls (e.g. common reference, common input, volume conduction or sample size problems). We do not include in Table 4 the connectivity toolboxes ibTB (Magri et al., 2009) and Toolconnect (Pastore et al., 2016), since they are not available under the links provided by the authors (accessed on 27.03.2019). We also do not list in Table 4 the following connectivity analysis toolboxes that are not focused on spike and LFP data analysis: Inform (Moore et al., 2017), HERMES (Niso et al., 2013), JIDT (Lizier, 2014), MVGC (Barnett and Seth, 2014), MuTe (Montalto et al., 2014), PyEntropy (Ince et al., 2009), SIFT (Delorme et al., 2011) and TrenTool (Lindner et al., 2011). TrenTool toolbox has a FieldTrip-compatible data structure.

**Table 4:** Comparison of connectivity analysis toolboxes for spike and LFP data. DTF – Directed Transfer Function (Kaminski and Blinowska, 1991), JPSTH – Joint Peri-Stimulus Time Histogram, MI – Mutual Information (Cover and Thomas, 2012), NC – Noise Correlations (Cohen and Kohn, 2011), PDC – Partial Directed Coherence (Baccalá and Sameshima, 2001), PPC – Pairwise Phase Consistency (Vinck et al., 2010), PSI – Phase Sloped Index (Nolte et al., 2004), RSEQ – statistical methods for detected Repeated SEQuences of synchronous spiking (Torre et al., 2016; Russo and Durstewitz, 2017; Staude et al., 2010; Quaglio et al., 2017), SFC – Spike-Field Coherence, STAT – STATistical tools, STD – Spike-Train Dissimilarity measures, STTC – Spike Time Tiling Coefficient (Cutts and Eglen, 2014), WPL – Weighted Phase Lag index (Vinck et al., 2011).

| Toolbox | (Cross)-<br>corre-<br>lation | Cohe-<br>rence | Granger<br>causality | Phase-amplitude<br>coupling | Phase-locking<br>value | Spike-triggered<br>average | Spike-field<br>cohe-<br>rence | Unique<br>features |
| --- | --- | --- | --- | --- | --- | --- | --- | --- |
| Brainstorm | + | + | + | + | + | + | – | STAT |
| Chronux | – | + | – | – | – | + | + | STAT |
| Elephant | + | + | – | – | – | + | + | RSEQ, STD,<br>STTC |
| FieldTrip | + | + | + | + | + | + | + | DTF, JPSTH, MI,<br>NC, PDC, PPC,<br>PSI, STAT, WPL |
| SPIKY | – | – | – | – | – | – | – | STD |

Compared to other toolboxes from Table 4,

- Brainstorm, Elephant and FieldTrip provide most versatile set of connectivity measures: while Field-Trip provides many classic and recent pairwise connectivity and synchronization measures, Elephant provides tools for multivariate analysis of high-order correlations in spike trains (see Subsections 4.1-4.2);
- Brainstorm tutorials for connectivity measures are actively developing ^39^; Chronux has examples for connectivity measures for real-world data in tutorial presentations; FieldTrip provides detailed tutorials on connectivity analysis for simulated and real-world data; Elephant provides examples for connectivity measures with simulated data;
- Chronux and FieldTrip compute confidence intervals for connectivity measures with jackknife re-sampling or variance estimates across trials, correspondingly (see Subsections 4.1-4.2); Brainstorm computes significance values for common connectivity measures, Elephant does not compute statistics on common connectivity measures.

To provide a better feeling of connectivity measures, we classify in Table 5 connectivity and synchronization measures mentioned in Table 4. We indicate for which signals the measure is applicable (Input), whether the measure is directed or not (Directed), is defined in time or frequency domain (Domain) and is bi- or multivariate (Dimension).

**Table 5:** Classification of synchronization and connectivity measures implemented in toolboxes listed in Table 4 regarding whether the measure is directed or not (Directed), is defined in time or frequency domain (Domain) and is bi- or multivariate (Dimension).

| Measure | Directed | Domain | Dimension |
| --- | --- | --- | --- |
| Correlation and cross-correlation (CC) | — | Time | Bivariate |
| Coherence | — | Frequency | Bivariate |
| Directed transfer information (DTF) | + | Frequency | Multivariate |
| Granger causality (GC) | + | Time, frequency | Bivariate |
| Imaginary part of coherency (iCOH) | — | Frequency | Bivariate |
| ISI and SPIKE distance, SPIKE synchronization (STD) | — | Time | Bivariate |
| Joint peri-stimulus time histogram (JPSTH) | — | Time | Bivariate |
| Mutual information (MI) | — | Time | Bivariate |
| Noise correlation (NC) | — | Time | Bivariate |
| Phase amplitude coupling (PAC) | — | Frequency | Bivariate |
| Partial coherence (pCOH) | + | Frequency | Bivariate |
| Partial directed coherence (pdCOH) | + | Frequency | Multivariate |
| Phase-locking value (PLV) | — | Frequency | Bivariate |
| Pairwise phase consistency (PPC) | + | Frequency | Bivariate |
| Phase slope index (PSI) | + | Frequency | Bivariate |
| statistical methods for detecting Repeated SEquences of synchronous spiking (RSEQ) | — | Time | Multivariate |
| Spike field coherence (SFC) | — | Time | Bivariate |
| van Rossum and Viktor-Purpura spike train dissimilarity measures (STD) | — | Time | Bivariate |
| Spike time tiling coefficient (STTC) | — | Time | Bivariate |
| Weighted phase lag index (WPL) | — | Frequency | Bivariate |

### 4.1 Comparing common tools: correlation, cross-correlation, coherence, Granger causality, phase-amplitude coupling, phase-locking value, spike-field coherence and spike-triggered average

In this subsection we compare implementations of common synchronization and connectivity measures for toolboxes from Table 4: correlation, cross-correlation, coherence, Granger causality, phase-amplitude coupling, phase-locking value, spike-field coherence and spike-triggered average.

Brainstorm and Elephant implement correlation, a pairwise non-directional time-domain connectivity measure. Brainstorm computes Pearson correlation coefficient (or optionally covariance) between spike trains and p-value of its significance; correlation is computed equivalently to MATLAB `corrcoef` function but in a faster vectorized way for *N* > 2 input signals. Elephant computes either Pearson correlation coefficient between binned spike trains (without additional statistics), pairwise covariances between binned spike trains (without additional statistics) or spike time tiling coefficient (STTC) introduced in (Cutts and Eglen, 2014). STTC, compared to correlation index introduced in (Wong et al., 1993), is described as not dependent on signals firing rate, correctly discriminating between lack of correlation and anti-correlation etc. (Cutts and Eglen, 2014). There is also a MATLAB STTC implementation^40^.

Cross-correlation is correlation between two signals computed for different time lags of one signal against the other. Elephant and FieldTrip implement cross-correlation, a pairwise non-directional time-domain connectivity measure. Between two binned spike trains Elephant computes cross-correlation for user-defined window with optional correction of border effect, kernel smoothing (for boxcar, Hamming, Hanning and Bartlett) and normalization. Between two LFP signals Elephant computes the standard unbiased estimator of the cross-correlation function (Stoica et al., 2005, Eq. 2.2.3) for user-defined time-lags without additional statistics across trials; note that biased estimator of the cross-correlation function is more accurate as discussed in (Stoica et al., 2005). FieldTrip computes cross-correlation between two spike channels for user-defined time lags and bin size (correlogram can optionally be debiased depending on data segment length). FieldTrip computes shuffled and unshuffled correlograms: if two channels are independent, the shuffled cross-correlogram should be the same as unshuffled.

Brainstorm, Chronux, Elephant and FieldTrip implement coherence, a frequency-domain equivalent of cross-correlation (Bastos and Schoffelen, 2016):

- Brainstorm implements coherence according to (Carter, 1987) computing also p-values of parametric significance estimation;
- Chronux computes coherence between two (binned) point-processes or LFP signals using multitaper methos, with confidence intervals or jackknife resampled error bars;
- Elephant computes coherence using Welch’s method with phase lags but without additional statistics. Computing coherence across trials is not supported in the considered version;
- FieldTrip computes coherence according to (Rosenberg et al., 1989) with variance estimate across trials. Additionally, FieldTrip provides computing of partial coherence according to (Rosenberg et al., 1998), partial directed coherence (Baccalá and Sameshima, 2001) and imaginary part of coherency (Nolte et al., 2004) with variance across trials. Partial directed coherence (PDC) is a directional measure. Compared to coherence, PDC is shown to reflect a frequency-domain representation of the concept of Granger causality (Baccalá and Sameshima, 2001).

Elephant does not provide built-in tools to compare coherence values between two conditions, Chronux provides a two-group test, FieldTrip provides an independent samples Z-statistic via `ft_freqstatistics` function by the method described in (Maris et al., 2007), Brainstorm is using FieldTrip `ft_freqstatistics` function.

Brainstorm and FieldTrip implement Geweke’s extension of the original time-domain concept of Granger causality (GC) introduced in (Granger, 1969) to the frequency domain (Geweke, 1982). GC implemented in Brainstorm and FieldTrip is a frequency-domain pairwise directional measure of connectivity. FieldTrip GC implementation is based on (Brovelli et al., 2004). The multivariate autoregressive (MVAR) model in FieldTrip uses biosig or BSMART toolboxes implementation on user choice, which are included in FieldTrip. FieldTrip computes variance of GC values across trials. Neither Brainstorm nor FieldTrip provide built-in tools/prescribed procedure to statistically compare GC values between conditions. Different to FieldTrip, Brainstorm computes as well time-resolved GC between two signals using two Wald statistics according to (Geweke, 1982) and (Hafner and Herwartz, 2008). The directed transfer function and partial directed coherence are multivariate extensions of Granger causality (Blinowska, 2011).

In Figure 6 we compare values of several connectivity measures computed in Brainstorm, Chronux and FieldTrip for simulated data with autoregressive models^41^ according to Eq. (5) (computing coherence across trials is not included in the considered Elephant version).

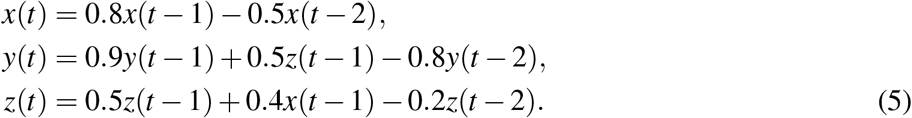

**Figure 6:**
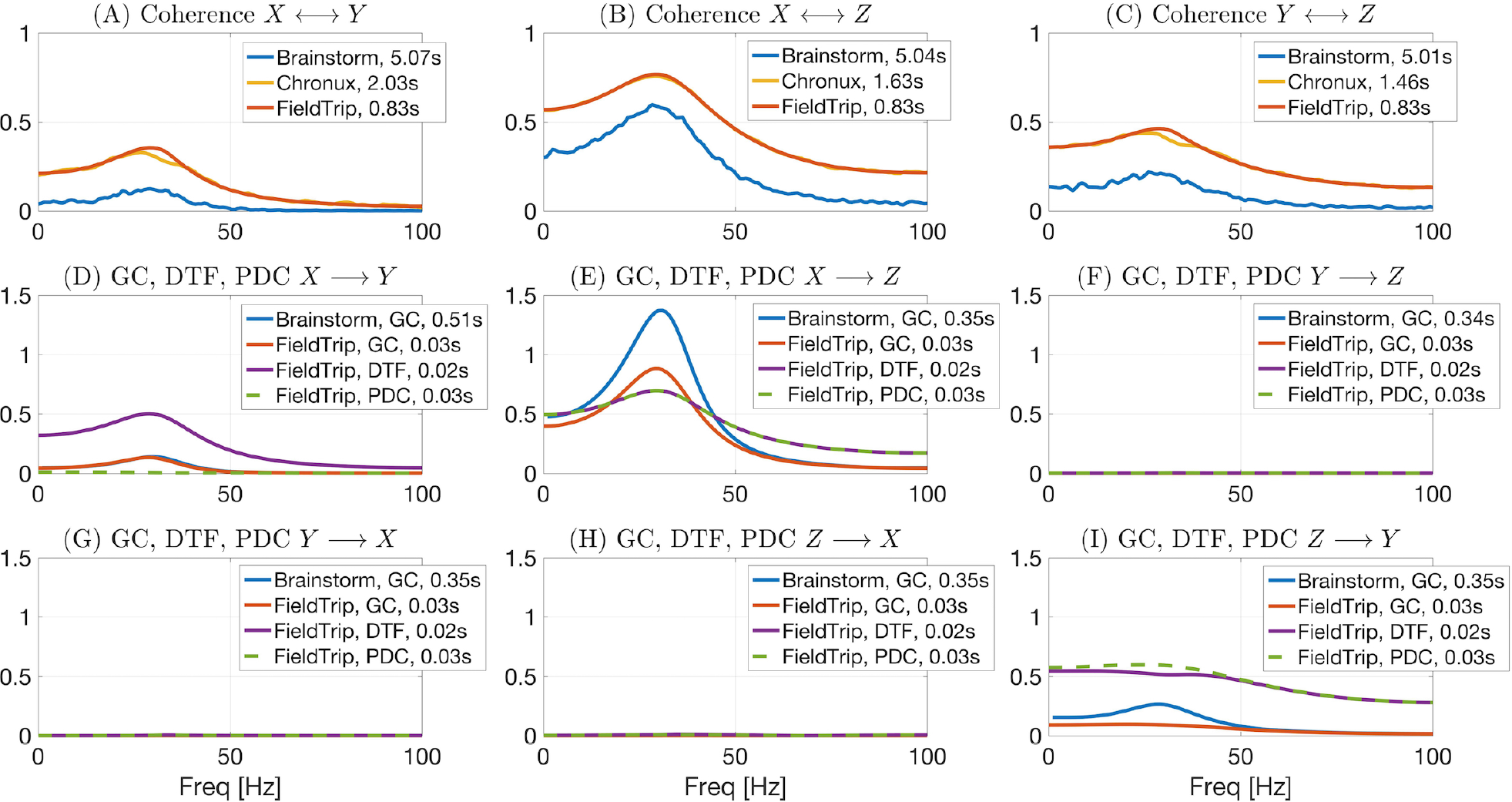
Comparing Brainstorm, Chronux and FieldTrip implementations of connectivity measures for signals simulated by autoregressive models (see Eq. (5)). While coherence is non-directional, Granger Causality (GC), Directed Transfer Function (DTF, see Subsection 4.2 for more details) and Partial directed Coherence (PDC) are directional measures. PDC allows to correctly detect interaction between signals (no direct *X → Y* interaction). Chronux and FieldTrip provide faster implementations compared to Brainstorm (see somputing times in plots legends) and return variance across trials. Brainstorm coherence values are noisier since there Welch method is used in contrast to multitapers (Chronux) or multivariate autoregressive modeling (FieldTrip).

Brainstorm and FieldTrip implement phase-amplitude coupling (PAC), a frequency-domain pairwise non-directional measure (Canolty et al., 2006; Samiee and Baillet, 2017; Voytek et al., 2010). FieldTrip implements two types of PAC^42^: mean vector length and modulation index according to (Tort et al., 2010). Brainstorm implements PAC according to (Özkurt and Schnitzler, 2011). Both Brainstorm and FieldTrip do not compute additional statistics on PAC.

Brainstorm and FieldTrip implement phase-locking value (PLV), a frequency-domain pairwise non-directional measure (Lachaux et al., 1999). PLV checks how consistent the phase relation between the two signals is across trials. We refer to (Vinck et al., 2011; Bastos and Schoffelen, 2016) for a comparison of different phase synchronization metrics and their biases. FieldTrip computes PLV based on (Lachaux et al., 1999) with a variance estimate using jackknife resampling.

Combination of spiking activity and LFP is often used to study rhythmic neuronal synchronization since spike-LFP measures are more sensitive than spike-spike synchronization measures (Vinck et al., 2012; Chakrabarti et al., 2014). To this end Brainstorm, Chronux, FieldTrip and Elephant implement a spike-field coherence (SFC), a frequency-domain pairwise non-directional measure. Brainstorm implements SFC according to (Fries et al., 2001) for user-defined window size around spikes without additional statistics computed. Chronux implements SFC with a multitaper approach for user-defined tapers and frequency band, computing also a confidence level of coherency and jackknife or standard error bars. FieldTrip computes SFC with variance across trials (see details in the corresponding tutorial^43^). Elephant implements SFC using standard Python `scipy.signal.coherence()` function, no additional statistics is computed.

One of the first steps in the analysis of spike-field coupling is computing of a spike-triggered average (STA) of LFP that is an average LFP voltage within a small window of the time around every spike. While neither Brainstorm nor Elephant compute any additional statistic on STA, Chronux computes STA with an optional kernel smoothing and calculates bootstrapped standard error on computed values and FieldTrip computes mean and variance of STA values.

### 4.2 Description of unique tools

In this subsection we describe unique tools of the toolboxes from Table 4. Elephant provides five recent statistical tools to study higher-order correlations and synchronous spiking events in parallel spike trains:

- ASSET (Analysis of Synchronous Spike EvenTs) implements the method from (Torre et al., 2016) and is an extension of the visualization method from (Schrader et al., 2008). ASSET assesses the statistical significance of simultaneous spike events (SSE) and aims to detect such events that cannot be explained on the basis of rate coding mechanisms and arise from spike correlations on shorter time scale;
- CAD (Cell Assembly Detection) implements the method from (Russo and Durstewitz, 2017) for capturing structures of higher-order correlations in massively parallel spike train recordings with arbitrary time lags and at multiple time-scale; CAD makes statistical parametric testing between each pair of neurons followed by an agglomerative recursive algorithm aiming to detect statistically precise repetitions of spikes in the data;
- CuBIC (Cumulant Based Inference of higher order Correlations) implements a statistical method (Staude et al., 2010) for detecting higher order correlations in parallel spike train recordings;
- SPADE (Spike Pattern Detection and Evaluation) implements the method from (Quaglio et al., 2017) for assessing the statistical significance of repeated occurrences of spike sequences (spatio-temporal patterns) based on recent methods in (Torre et al., 2013; Quaglio et al., 2017). SPADE aims to overcome computational and statistical limits in detecting repeated spatio-temporal patterns within massively parallel spike trains (Quaglio et al., 2017), see (Quaglio et al., 2018) for a recent review of methods for identification of spike patterns in massively parallel spike trains;
- UE (Unitary Event analysis) implements the statistical method from (Grün et al., 1999, 2002) for analyzing excess spike correlations between simultaneously recorded neurons. This method compares the empirical spike coincidences to the expected number on the basis of firing rate of the neurons.

Elephant and SPIKY toolboxes allow to compute measures of spike train dissimilarity (also referred as measures of spike train synchrony). Elephant implements well-known time-scale dependent van Rossum (van Rossum, 2001) and (Victor and Purpura, 1996) dissimilarity distances whereas SPIKY implements three recent parameter-free time-scale independent measures: ISI-distance (Kreuz et al., 2007), SPIKY distance (Kreuz et al., 2012) and SPIKE synchronization (Quiroga et al., 2002). We refer to (Chicharro et al., 2011; Kreuz et al., 2012; Mulansky et al., 2015) for a comparison of dissimilarity measures. Note also MATLAB implementations of dissimilarity measures at J.D. Victor^44^ and T. Kreuz^45^ web-sites.

FieldTrip, compared to other toolboxes from Table 4, computes and visualizes^46^ the following classic and recent connectivity and synchronization measures:

- directed transfer function (DTF) introduced in (Kaminski and Blinowska, 1991) is a multivariate frequency-domain directional connectivity measure; FieldTrip computes it according to (Kaminski and Blinowska, 1991) from cross-spectral density with a variance across trials. DTF, compared to GC, makes a multivariate spectral decomposition, the advantage of this approach is that interaction between all channels is taken into account (see, e.g., Figure 6 in Subsection 4.1). However pairwise measures yield more stable results since they involve fitting fewer parameters (Blinowska, 2011; Bastos and Schoffelen, 2016);
- joint peri-stimulus time histogram (JPSTH) is a pairwise time-domain non-directional measure between spike trains that allows to gain insight into temporal evolution of spike-spike correlations (Brown et al., 2004; Aertsen et al., 1987). To check whether the resulted JPSTH is caused by task-induced fluctuations of firing rate or by temporal coordination not time-locked to stimulus onset, FieldTrip also computes JPSTH with shuffling subsequent trials. We illustrate JPSTH visualization with FieldTrip tools in Figure 7;
- mutual information (MI) is a pairwise time-domain non-directional connectivity measure. FieldTrip computes MI using implementation from ibtb toolbox (Magri et al., 2009) without additional statistics;
- noise correlations (NC) is a non-directional pairwise time-domain measure that can be computed between two spike trains; NC measures whether neurons share trial-by-trial fluctuations in their firing rate; different to so called signal correlations (SC), these fluctuations are measured over repetitions of identical experimental conditions, i.e. are not driven by variable sensory or behaviorally conditions;
- phase-coupling pairwise spike-field measures compute the phases of spikes relative to the ongoing LFP with a discrete Fourier transform of an LFP segment around the spike time (Vinck et al., 2012). FieldTrip implements recent methods from (Vinck et al., 2012): angular mean of spike phases, Rayleigh p-value and pairwise-phase consistency according to the method in (Vinck et al., 2010). We refer to (Vinck et al., 2010; Bastos and Schoffelen, 2016) for a discussion and comparison of these measures;
- phase-slope index (PSI) is a directional pairwise frequency-domain measure that can be computed between two signals from their complex-valued coherency. FieldTrip computes PSI according to (Nolte et al., 2008) with variance across trials;
- pairwise phase consistency (PPC) is a directional pairwise frequency-domain measure that can be computed from the distribution of pairwise differences of the relative phases. PPC compared to PLV is not biased by sample size (Bastos and Schoffelen, 2016). FieldTrip computes PPC with leave-one-out variance estimate;
- weighted phase-lag index (WPL) introduced in (Vinck et al., 2011) is a non-directional pairwise frequency-domain measure computed from cross-spectral density between two signals. WPL was introduced to solve the problem with sensitivity of phase-lag index (Stam et al., 2007) to volume-conduction and noise (Vinck et al., 2011). FieldTrip computes WPL according to (Vinck et al., 2011) with variance across trials.

**Figure 7:**
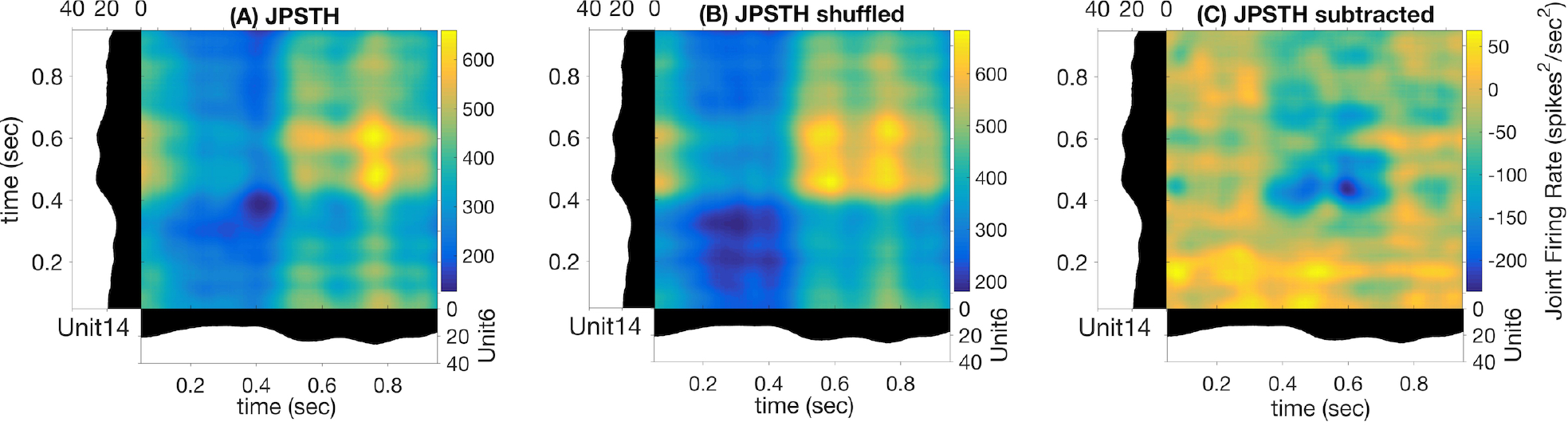
Illustration of FieldTrip functionality: joint peri-stimulus time histogram (JPSTH) (A), shuffled JPSTH (B) and their difference (C) for test dataset between M1 units 6 and 42, monkey MM (colorbar values range is different for each subplot, this range is not adjustable outside of FieldTrip).

## 5 SPECIALIZED TOOLBOXES FOR DIMENSIONALITY REDUCTION AND GENERALIZED LINEAR MODELING

In this section we overview specialized toolboxes for dimensionality reduction (Subsection 5.1) and generalized linear modeling (Subsection 5.2). Compared to Table 1, we do not provide in the corresponding tables for specialized toolboxes information on

- Import/Export since none of the considered toolboxes supports importing/exporting from specialized spike data formats;
- GUI since only DataHigh toolbox provides GUI (see details below).

### 5.1 Toolboxes for dimensionality reduction

Dimensionality reduction of neural data allows to obtain a simplified low-dimensional representation of neural activity. In Table 6 we compare open-source toolboxes for dimensionality reduction of neural data (note also a list of dimensionality reduction software actively updating at B. Yu web-site^47^). See examples for application of DataHigh, dPCA and TCA toolboxes in our open MATLAB script.

**Table 6:**
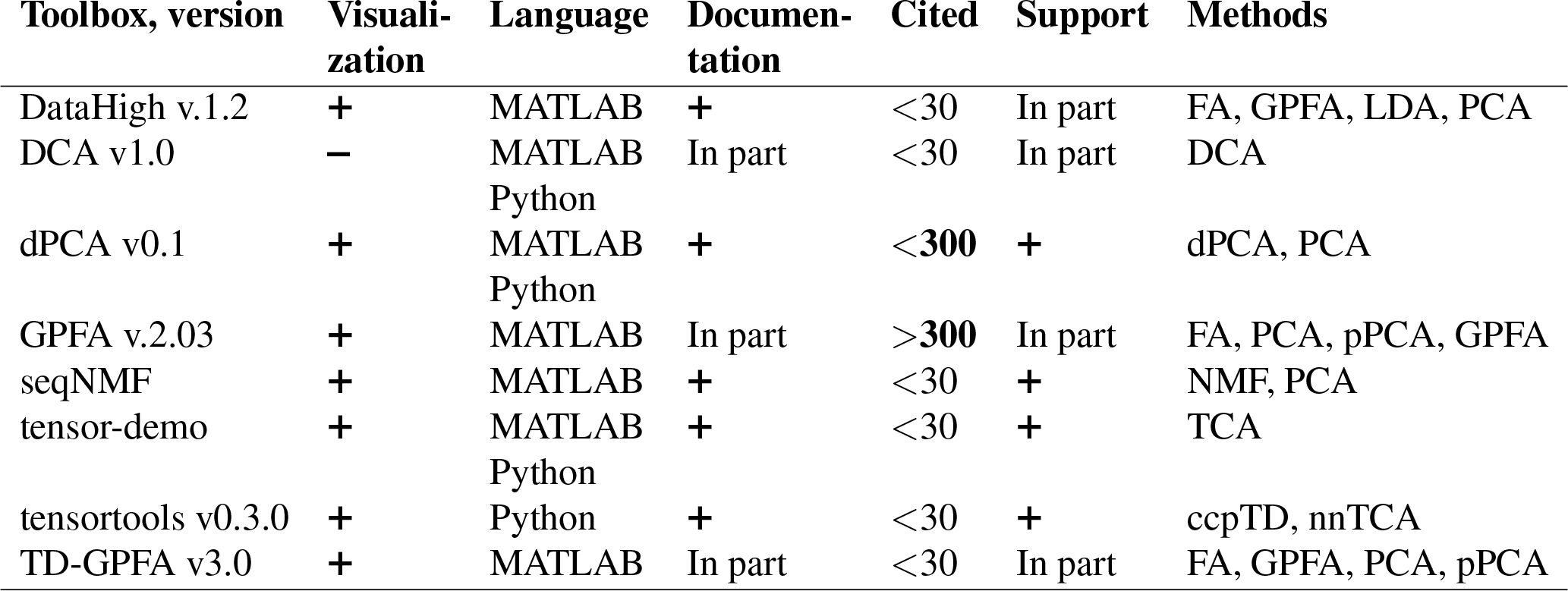
Features of open-source dimensionality reduction toolboxes regarding visualization tools, principal and usage programming language, availability of documentation, number of citations, and support by updates at least once per year. ccpTD – coupled canonical polyadic Tensor Decomposition, DCA – Distance Covariances Analysis, (GP)FA – (Gaussian Process) Factor Analysis, LDA – Fisher’s Linear Discriminant Analysis, NMF – Non-negative Matrix Factorization, (d,p)PCA – (demixed, probabilistic) Principal Component Analysis, (nn)TCA – (non-negative) Tensor Component Analysis

We have indicated “In part” in Documentation column for GPFA and TD-GPFA toolboxes since they provide usage examples and readme files with notes on parameters choice but but neither detailed manual nor tutorial, they refer to the original publication (Yu et al., 2009) for details. We have indicated “In part” in Documentation column for DCA tool since it provides neither manual nor tutorial (only example of use in MATLAB script comments). DataHigh and GPFA toolboxes are not uploaded to GitHub or any other public version control system preventing from tracking version changes and submitting bugs. DCA and TD-GPFA toolboxes have not been updated during the last 2 years.

Compared to other toolboxes from Table 6,

- DataHigh provides a user-friendly GUI illustrating algorithm steps such as choice of bin size, smoothing, components number etc.;
- dPCA is applied on trial-averaged spiking activity; dPCA breaks down the neural activity into components each of which relates to time (condition-independent component) or a single experimental condition of the task; the idea is an easier task-relevant interpretation compared to the standard PCA or ICA; the results can be summarized in a single figure (Kobak et al., 2016);
- TD-GPFA allows to extract low-dimensional latent structure from time series in the presence of delays;
- tensor-demo and tensortools allow to reduce dimensionality both across and within trials (Williams et al., 2018).

In Table 7 we outline additional dimensionality reduction tools provided by the toolboxes.

**Table 7:** Comparing dimensionality reduction toolboxes: diagnostic and statistical tools. In Statistical tests column we indicate whether the toolbox provides possibility to measure significance of results and provides permutation or re-shuffling tests on the data.

| Toolbox | Cross-validation | Tool to select optimal dimensions number | Fitting error, variance explained | Statistical tests |
| --- | --- | --- | --- | --- |
| DataHigh | + | + | + | — |
| DCA | — | — | — | — |
| dPCA | + | + | + | + |
| GPFA | + | + | + | — |
| seqNMF | + | + | + | + |
| tensor-demo | — | + | + | — |
| tensortools | — | + | + | + |
| TD-GPFA | + | + | + | — |

It is important to check whether input data fit model assumptions when applying dimensionality reduction methods: whether the data are allowed to be non-stationary, contain outliers, observational noise or be correlated, whether recorded activity evolves in a low-dimensional manifold, which sample size is sufficient etc. Discussing model assumptions for each of dimensionality reduction methods is beyond the scope of this paper, we refer to the original papers and to the model assumptions for applying principal component analysis (PCA) formulated in (Shlens, 2014).

### 5.2 Toolboxes for GLM analysis

Generalized linear models (GLMs) are often applied for predicting spike counts with the aim to understand which factors influence simultaneous spiking activity: whether it is predicted by the past or concurrent neural activity of the same or remote brain area or by external covariates. In Table 8 we overview major open-source toolboxes for GLM analysis. These toolboxes do not contain any general spike data analysis functions besides GLM analysis since they are either GLM tutorials or codes related to particular analysis made in the paper.

**Table 8:** Features of open-source toolboxes for generalized linear modeling of spike data regarding visualization tools, principal and usage programming language, availability of documentation, number of citations (for the paper with the introduced method), support by updates at least once per year and implemented methods. cGLM – GLM with coupling filters, gGLM – linear Gaussian GLM, GNM – Generalized Nonlinear Model (Butts et al., 2011), GQM – Generalized Quadratic Model (Park and Pillow, 2011), pGLM – Poisson GLM (Truccolo et al., 2005), ppGLM – point-process GLM (Paninski et al., 2007), NIM – Nonlinear Input Model (McFarland et al., 2013), SHF – Spikes and covariates History Filters, STB – Smooth Temporal Basis

| Toolbox, version | Methods | Visuali-<br>zation | Language | Documen-<br>tation | Cited | Support |
| --- | --- | --- | --- | --- | --- | --- |
| Case-Studies | pGLM | + | MATLAB | + | <30 | + |
| GLMcode1 | pGLM | + | MATLAB | + | <30 | — |
| GLMcode2 | pGLM | + | MATLAB | + | <30 | — |
| GLMspiketools v1 | cGLM, pGLM, SHF, STB | + | MATLAB | + | >900 | + |
| GLMspike-traintutorial | cGLM, gGLM, pGLM, SHF | + | MATLAB | + | >900 | + |
| neuroGLM | pGLM, SHF, STB | + | MATLAB | + | >90 | + |
| NIMclass v1.0 | GLM, GQM, GNM, NIM | — | MATLAB | — | >90 | + |
| nStat v2 | ppGLM | + | MATLAB | In part | <30 | + |
| spykesML v0.1.dev | pGLM, SHF | — | Python | + | <30 | + |

GLMcode1 and GLMcode2 codes are not uploaded to GitHub or any other version control system as they implement methods for particular analysis made in the papers (see below) are not supposed to be updated.

Note that

- Case-Studies implements (see folders Chapter 9, 10, 11 on GitHub^48^) basic steps of Poisson GLM fitting with history dependence to the data on sample datasets for the corresponding book (Kramer and Eden, 2016);
- GLMcode1, GLMcode2 implement the code for the papers (Glaser et al., 2018) and (Lawlor et al., 2018);
- examples of use for nStat toolbox are located in `helpfiles` folder in the corresponding GitHub repository;
- spykesML tool provides comparison of GLM performance with several methods from modern machine learning approaches (including neural networks);
- NIMclass uses MATLAB optimization toolbox and contains many examples for real-world data;
- GLMspiketraintutorial is a tutorial for teaching purposes. It is not memory-efficient implemented, but it makes easy to understand the basic steps of Poisson and Gaussian GLMs fitting, analysis and comparison for spike data ^49^. neuroGLM and GLMspiketools are more advanced tools with efficient memory implementation. Additionally to GLMspiketraintutorial, they support some advanced GLM features such as smooth temporal basis functions for spike-history filters, different time-scales for stimulus and spike-history components etc.

## 6 CONCLUSIONS

In this review we have compared major open-source toolboxes for spike and local field potentials (LFP) processing and analysis. We have compared toolboxes functionality, statistical and visualization tools, documentation and support quality. Besides summarizing information about toolboxes in comparison tables, we have discussed and illustrated particular toolboxes functionality and implementations, also in our open MATLAB code. Below we summarize the comparisons that we made for general spike and LFP analysis toolboxes and toolboxes with connectivity tools.

Each considered toolbox has its own advantages:

- Brainstorm: graphical user interface (GUI), versatile and cross-checked functionality (highly-cited), statistical tools, detailed tutorials with recommendations on parameters choice, support of many file formats, active user discussion community and regular hands-on sessions, fast Morlet wavelet transform implementation;
- Chronux: versatile and cross-checked functionality (highly-cited), statistical tools (measures of variance across trials and statistical comparing between different conditions), detailed documentation, convenient data analysis pipeline for programming-oriented users (detailed code comments and modular code design);
- Elephant: support of many file formats, versatile functionality with implementation of classic and recent methods for spike-spike connectivity and synchronization analysis, fast Morlet wavelet transform implementation;
- FieldTrip: versatile and cross-checked functionality (highly-cited), statistical tools (measures of variance across trials and statistical comparing between different conditions), detailed tutorials with recommendations on parameters choice, support of many file formats, active user discussion community and regular hands-on sessions, flexible visualization tools, convenient data analysis pipeline for programming-oriented users (detailed code comments and modular code design), versatile filtering, connectivity and synchronization analysis tools, fast and accurate line noise removal;
- gramm: quick publication-quality PSTH, raster plots and tuning curves with many easily adjustable plot properties;
- Spike Viewer: GUI, support of many file formats;
- SPIKY: GUI, implementation of recent spike train dissimilarity measures.

## 7 LIST OF TOOLBOXES AND TOOLS IN ALPHABETICAL ORDER WITH LINKS

Below all the considered toolboxes are provided with a brief description, reference to the paper where the toolbox was introduced and a link for downloading.

- Brainstorm^50,51^ (Tadel et al., 2011) – a MATLAB toolbox for the analysis of brain recordings: MEG, EEG, fNIRS, ECoG, depth electrodes and animal invasive neurophysiology;
- BSMART^52^ (Brain-System for Multivariate AutoRegressive Time series) (Cui et al., 2008) – a MATLAB/C toolbox for spectral analysis of continuous neural data recorded from several sensors;
- Case-Studies^53^ – a MATLAB set of examples on sample datasets accompanying the corresponding book (Kramer and Eden, 2016);
- Chronux^54^ (Bokil et al., 2010) – a MATLAB package for the analysis of neural data;
- DataHigh^55^ (Cowley et al., 2013) – a MATLAB-based graphical user interface to visualize and interact with high-dimensional neural population activity;
- DATA-MEAns^56^ (Bonomini et al., 2005) – a Delphi7 tool for the classification and management of neural ensemble recordings;
- DCA^57^ (Cowley et al., 2017) (distance covariance analysis) – an implementation (MATLAB and Python) of the linear dimensionality reduction method that can identify linear and nonlinear relation-ships between multiple datasets;
- dPCA^58^ (demixed Principal Component Analysis) (Kobak et al., 2016) – a MATLAB implementation of the linear dimensionality reduction technique that automatically discovers and highlights the essential features of complex population activities;
- Elephant^59,60^ (Yegenoglu et al., 2017) – an Electrophysiology Analysis Toolkit in Python. Elephant toolbox includes functionality from earlier developed toolboxes CSDPlotter^61^ (Pettersen et al., 2006) and iCSD 2D^62^, it is a direct successor of NeuroTools;
- FieldTrip^63,64^ (Oostenveld et al., 2011) – a MATLAB toolbox for advanced analysis of MEG, EEG, and invasive electrophysiological (spike and LFP) data;
- FIND^65^ (Meier et al., 2008) – a MATLAB toolbox for the analysis of neuronal activity;
- GLMcode1 – a MATLAB code implementing data analysis for particular publication (Glaser et al., 2018) with GLM fitting to analyze factors contributing to neural activity (this code is available from the authors upon request);
- GLMcode2^66^ (Perich et al., 2018) – a MATLAB code implementing data analysis for particular publication (Lawlor et al., 2018) with GLM fitting to estimate preferred direction for each neuron;
- GLMspikestools^67^ (Pillow et al., 2008) – a Generalized Linear Modeling tool for single and multi-neuron spike trains;
- GLMspiketraintutorial^68^ (Pillow et al., 2008) – a simple tutorial on Gaussian and Poisson GLMs for single and multi-neuron spike train data;
- GPFA^69^ (Gaussian-Process Factor Analysis) (Yu et al., 2009) – a MATLAB implementation of the method extracting low-dimensional latent trajectories from noisy, high-dimensional time series data. It combines linear dimensionality reduction (factor analysis) with Gaussian-process temporal smoothing in a unified probabilistic framework;
- gramm^70,71^ (Morel, 2018) – a plotting MATLAB toolbox for quick creation of complex publication-quality figures;
- HERMES^72^ (Niso et al., 2013) – a MATLAB toolbox for assessing conectivity and sinchronization between time series;
- ibTB^73^ (Information Breakdown Toolbox) (Magri et al., 2009) – a C/MATLAB toolbox for fast information analysis of multiple-site LFP, EEG and spike train recordings;
- Inform^74^ (Moore et al., 2017) – a cross-platform C library for information analysis of dynamical systems;
- infoToolbox^75^ (Magri et al., 2009) – a toolbox for the fast analysis of multiple-site LFP, EEG and spike train recordings;
- JIDT^76^ (Lizier, 2014) – an information-theoretic Java toolbox for studying dynamics of complex systems;
- MEAbench^77^ (Wagenaar et al., 2005) – a C++ toolbox for multi-electrode data acquisition and online analysis;
- MEA-tools^78^ (Egert et al., 2002) – a collection of MATLAB-based tools to analyze spike and LFP data from extracellular recordings with multi-electrode arrays;
- MuTe^79^ (Montalto et al., 2014) – a MATLAB toolbox to compare established and novel estimators of the multivariate transfer entropy;
- MVGC^80^ (Multivariate Granger Causality MATLAB Toolbox) (Barnett and Seth, 2014) – a MATLAB toolbox facilitating Granger-causal analysis with multivariate multi-trial time series data;
- neuroGLM^81^ (Park et al., 2014) – an MATLAB tool, an extension of GLMspiketraintutorial allowing more advanced features of GLM modeling such as smooth basis functions for spike-history filters, memory-efficient temporal convolutions, different timescales for stimulus and spike-history components, low-rank parametrization of spatio-temporal filters, flexible handling of trial-based data;
- NIMclass^82,83^ (McFarland et al., 2013) – a MATLAB implementation of the nonlinear input model. In this model, the predicted firing rate is given as a sum over nonlinear inputs followed by a “spiking nonlinearity” function;
- nStat^84^ (neural Spike Train Analysis Toolbox) (Cajigas et al., 2012) – an object-oriented MATLAB toolbox that implements several models and algorithms for neural spike train analysis;
- OpenElectrophy^85,86^ (Garcia and Fourcaud-Trocmé, 2009) – a Python framework for analysis of intro- and exrta-cellular recordings;
- PyEntropy^87^ (Ince et al., 2009) – a Python module for estimating entropy and information theoretic quantities using a range of bias correction methods;
- seqNMF^88^ (Mackevicius et al., 2019) – a MATLAB toolbox for unsupervised discovery of temporal sequences in high-dimensional datasets with applications to neuroscience;
- SIFT^89^ (Delorme et al., 2011; Mullen, 2014) – a Source Information Flow MATLAB EEGLAB-compatible toolbox for analysis and visualization of multivariate causality and information flow between sources of electrophysiological (EEG/ECoG/MEG) activity;
- SigMate^90^ (Mahmud et al., 2012) – a MATLAB toolbox for extracellular neuronal signal analysis;
- sigTOOL^91^ (Lidierth, 2009) – a MATLAB toolbox for spike data analysis;
- Spike Viewer^92^ (Pröpper and Obermayer, 2013) – a multi-platform GUI application for navigating, analyzing and visualizing electrophisiological datasets;
- SPIKY^93,94^ (Kreuz et al., 2015) – a MATLAB graphical user interface that facilitates application of time-resolved measures of spike-train synchrony to both simulated and real data;
- SPKTool^95^ (Liu et al., 2011) – a MATLAB toolbox for spikes detection, sorting and analysis;
- spykesML^96^ (Benjamin et al., 2018) – a Python toolbox with a tutorial for comparing performance of GLM with modern machine-learning methods (neural networks, random forest etc.);
- STAR^97^ (Spike Train Analysis with R) (Pouzat and Chaffiol, 2009) – an R package to analyze spike trains;
- STAToolkit^98^ (Spike Train Analysis Toolkit) (Goldberg et al., 2009) – a MATLAB package for the information theoretic analysis of spike train data;
- tensor-demo^99^ – a MATLAB and Python package (available for both languages) for fitting and visualizing canonical polyadic tensor decompositions of higher-order data arrays;
- tensortools^100^ – a Python package for fitting and visualizing canonical polyadic tensor decompositions of higher-order data arrays;
- TD-GPFA^101^ (time-delayed Gaussian-Process Factor Analysis) (Lakshmanan et al., 2015) – a MATLAB implementation of GPFA method extension that allows for a time delay between each latent variable and each neuron;
- ToolConnect^102^ (Pastore et al., 2016) – a functional connectivity C# toolbox with GUI for in vitro networks;
- Trentool^103^ (Lindner et al., 2011) – a MATLAB toolbox for the analysis of information transfer in time series data. Trentool provides user friendly routines for the estimation and statistical testing of transfer entropy in time series data.

## CONFLICT OF INTEREST STATEMENT

The authors have declared that no competing interests exist.

## AUTHOR CONTRIBUTIONS

VU performed the reported study. VU wrote and AG edited the paper. Both authors have seen and approved the final manuscript.

## FUNDING

The work was supported by grants of the Deutsche Forschungsgemeinschaft through the Collaborative Research Center 889 “Cellular Mechanisms of Sensory Processing” and the Research Unit 1847 “The Physiology of Distributed Computing Underlying Higher Brain Functions in Non-Human Primates”, and by the European Commission through the H2020 project Plan4Act (FETPROACT-16 732266), all granted to AG. The funder had no role in study design, data collection and analysis, decision to publish, or preparation of the manuscript.

## DATA AVAILABILITY STATEMENT

The datasets analyzed and generated for this study can be found at (Perich et al., 2018; Lowet et al., 2015) and on GitHub^104^, correspondingly.

## Footnotes

1 https://github.com/ValentinaUn/Testing-open-source-toolboxes

2 according to Google Scholar in March 2019 (https://scholar.google.com)

3 when the code is available under a license which allows free redistribution and the creation of derived works

4 https://www.mathworks.com

5 https://www.python.org

6 https://github.com/OpenElectrophy/OpenElectrophy

7 https://github.com/piermorel/gramm/blob/master/gramm%20cheat%20sheet.pdf

8 https://elephant.readthedocs.io/en/latest/tutorial.html

9 https://github.com

10 https://github.com/NeuralEnsemble/python-neo

11 http://neuralensemble.org/neo/

12 see https://neuroimage.usc.edu/brainstorm/Introduction, Chronux folder dataio and http://www.fieldtriptoolbox.org/dataformat for details, correspondingly

13 One can recompile locfit by running `locfit/source/compile.m`

14 https://simonster.github.io/SpikeSortingSoftware/

15 https://www.cnsorg.org/software

16 https://grey.colorado.edu/emergent/index.php/Comparison_of_Neural_Network_Simulators

17 https://neuroimage.usc.edu/forums/

18 http://www.fieldtriptoolbox.org/discussion_list/

19 http://www.fieldtriptoolbox.org/workshop/

20 https://neuroimage.usc.edu/brainstorm/Training

21 https://neuroimage.usc.edu/brainstorm/Tutorials/Statistics

22 https://neuroimage.usc.edu/brainstorm/Tutorials/Statistics

23 https://neuroimage.usc.edu/brainstorm/e-phys/functions

24 https://spyke-viewer.readthedocs.io/en/latest/

25 http://www.fieldtriptoolbox.org/reference/ft_timelockstatistics/

26 https://github.com/piermorel/gramm/blob/master/gramm

27 https://neuroimage.usc.edu/brainstorm/e-phys/SpikeSorting?highlight=%28sorting%29

28 https://neuroimage.usc.edu/brainstorm/e-phys/functions

29 https://neuroimage.usc.edu/brainstorm/Tutorials/ArtifactsFilter

30 https://neuroimage.usc.edu/brainstorm/Tutorials/TimeFrequency

31 http://www.fieldtriptoolbox.org/example/determine_the_filter_characteristics/

32 http://www.fieldtriptoolbox.org/tutorial/timefrequencyanalysis/

33 http://chronux.org

34 http://www.fieldtriptoolbox.org/reference/ft_preprocessing

35 https://www.mathworks.com/help/signal/ug/remove-the-60-hz-hum-from-a-signal.html

36 http://chronux.org/chronuxFiles//Documentation/chronux/spectral_analysis/continuous/locdetrend.html

37 http://www.fieldtriptoolbox.org/reference/ft_preproc_detrend/

38 https://de.mathworks.com/help/matlab/data_analysis/detrending-data.html

39 https://neuroimage.usc.edu/brainstorm/Tutorials/Connectivity

40 https://github.com/Leo-GG/NeuroFun/blob/master/%2Bcorrel/calcSTTC.m

41 http://www.fieldtriptoolbox.org/tutorial/connectivity/

42 http://www.fieldtriptoolbox.org/reference/ft_crossfrequencyanalysis/

43 http://www.fieldtriptoolbox.org/tutorial/spikefield/

44 http://www-users.med.cornell.edu/~jdvicto/pubalgor.html

45 http://wwwold.fi.isc.cnr.it/users/thomas.kreuz/Source-Code/VanRossum.html

46 http://www.fieldtriptoolbox.org/reference/ft_connectivityplot/

47 http://users.ece.cmu.edu/~byronyu/software.shtml

48 https://github.com/Mark-Kramer/Case-Studies-Kramer-Eden

49 https://github.com/pillowlab/GLMspiketraintutorial

50 https://neuroimage.usc.edu/brainstorm/Introduction

51 https://github.com/brainstorm-tools/brainstorm3

52 http://www.brain-smart.org

53 https://github.com/Mark-Kramer/Case-Studies-Kramer-Eden

54 http://chronux.org

55 http://users.ece.cmu.edu/~byronyu/software/DataHigh/datahigh.html

56 http://cortivis.umh.es

57 https://github.com/BenjoCowley/dca

58 https://github.com/machenslab/dPCA

59 http://neuralensemble.org/elephant/

60 https://github.com/NeuralEnsemble/elephant/commits/master

61 https://github.com/espenhgn/CSDplotter

62 http://www.neuroinf.pl/Members/szleski/csd2d/toolbox

63 http://www.fieldtriptoolbox.org

64 https://github.com/fieldtrip/fieldtrip

65 http://find.bccn.uni-freiburg.de

66 https://crcns.org/data-sets/motor-cortex/pmd-1/about-pmd-1

67 http://pillowlab.princeton.edu/code_GLM.html

68 https://github.com/pillowlab/GLMspiketraintutorial

69 http://users.ece.cmu.edu/~byronyu/software.shtml

70 https://www.mathworks.com/MATLABcentral/fileexchange/54465-gramm-complete-data-visualization-toolbox-ggplot2-r-like

71 https://github.com/piermorel/gramm

72 http://hermes.ctb.upm.es

73 http://www.ibtb.org

74 https://github.com/ELIFE-ASU/Inform

75 http://www.infotoolbox.org

76 https://github.com/jlizier/jidt

77 http://www.danielwagenaar.net/meabench.html

78 http://material.brainworks.uni-freiburg.de/research/meatools/

79 https://figshare.com/articles/MuTE_toolbox_to_evaluate_Multivariate_Transfer_Entropy/1005245

80 http://www.sussex.ac.uk/sackler/mvgc/

81 https://github.com/pillowlab/neuroGLM

82 http://neurotheory.umd.edu/nimcode

83 https://github.com/dbutts/NIMclass

84 https://github.com/iahncajigas/nSTAT

85 http://neuralensemble.org/OpenElectrophy/

86 https://github.com/OpenElectrophy/OpenElectrophy

87 https://github.com/robince/pyentropy

88 https://github.com/FeeLab/seqNMF

89 https://sccn.ucsd.edu/wiki/SIFT

90 https://sites.google.com/site/muftimahmud/codes

91 http://sigtool.sourceforge.net/

92 https://github.com/rproepp/spykeviewer

93 https://github.com/mariomulansky/PySpike

94 http://wwwold.fi.isc.cnr.it/users/thomas.kreuz/Source-Code/SPIKY.html

95 https://sourceforge.net/projects/spktool/files/latest/download

96 https://github.com/KordingLab/spykesML

97 https://sites.google.com/site/spiketrainanalysiswithr/

98 https://omictools.com/statoolkit-tool

99 https://github.com/ahwillia/tensor-demo

100 https://github.com/ahwillia/tensortools

101 https://github.com/karts25/NeuralTraj

102 http://software.incf.org/software/toolconnect

103 https://trentool.github.io/TRENTOOL3/

104 https://github.com/ValentinaUn/Testing-open-source-toolboxes

## References

Abramson, I. (1982). On bandwidth variation in kernel estimates-a square root law. The annals of Statistics, 1217–1223

Aertsen, A., Bonhoeffer, T., and Krüger, J. (1987). Coherent activity in neuronal populations: analysis and interpretation. Physics of cognitive processes, 1–34 doi:10.1523/JNEUROSCI.23-17-06798.2003

Baccalá, L. and Sameshima, K. (2001). Partial directed coherence: a new concept in neural structure determination. Biological cybernetics 84, 463–474. doi:10.1007/PL00007990

Barnett, L. and Seth, A. (2014). The MVGC multivariate Granger causality toolbox: a new approach to Granger-causal inference. Journal of neuroscience methods 223, 50–68. doi:10.1016/j.jneumeth.2013.10.018

Bastos, A. and Schoffelen, J.-M. (2016). A tutorial review of functional connectivity analysis methods and their interpretational pitfalls. Frontiers in systems neuroscience 9, 175. doi:10.3389/fnsys.2015.00175

Benjamin, A., Fernandes, H., Tomlinson, T., Ramkumar, P., VerSteeg, C., Chowdhury, R., et al. (2018). Modern machine learning as a benchmark for fitting neural responses. Frontiers in computational neuroscience 12. doi:10.3389/fncom.2018.00056

Blinowska, K. (2011). Review of the methods of determination of directed connectivity from multichannel data. Medical & biological engineering & computing 49, 521–529. doi:10.1007/s11517-011-0739-x

Bokil, H., Andrews, P., Kulkarni, J., Mehta, S., and Mitra, P. (2010). Chronux: a platform for analyzing neural signals. Journal of neuroscience methods 192, 146–151. doi:10.1016/j.jneumeth.2010.06.020

Bonomini, M., Ferrandez, J., Bolea, J., and Fernandez, E. (2005). DATA-MEAns: an open source tool for the classification and management of neural ensemble recordings. Journal of neuroscience methods 148, 137–146. doi:10.1016/j.jneumeth.2005.04.008

Brovelli, A., Ding, M., Ledberg, A., Chen, Y., Nakamura, R., and Bressler, S. (2004). Beta oscillations in a large-scale sensorimotor cortical network: directional influences revealed by granger causality. Proceedings of the National Academy of Sciences 101, 9849–9854. doi:10.1073/pnas.0308538101

Brown, E., Kass, R., and Mitra, P. (2004). Multiple neural spike train data analysis: state-of-the-art and future challenges. Nature neuroscience 7, 456. doi:10.1038/nn1228

Bruns, A. (2004). Fourier-, Hilbert-and wavelet-based signal analysis: are they really different approaches? Journal of neuroscience methods 137, 321–332

Butts, D., Weng, C., Jin, J., Alonso, J.-M., and Paninski, L. (2011). Temporal precision in the visual pathway through the interplay of excitation and stimulus-driven suppression. Journal of Neuroscience 31, 11313–11327. doi:10.1523/JNEUROSCI.0434-11.2011

Cajigas, I., Malik, W., and Brown, E. (2012). nSTAT: open-source neural spike train analysis toolbox for Matlab. Journal of neuroscience methods 211, 245–264. doi:10.1016/j.jneumeth.2012.08.009

Canolty, R., Edwards, E., Dalal, S., Soltani, M., Nagarajan, S., and Kirsch, H. e. a. (2006). High gamma power is phase-locked to theta oscillations in human neocortex. science 313, 1626–1628. doi:10.1126/science.1128115

Carter, G. (1987). Coherence and time delay estimation. Proceedings of the IEEE 75, 236–255. doi: 10.1109/PROC.1987.13723

Chakrabarti, S., Martinez-Vazquez, P., and Gail, A. (2014). Synchronization patterns suggest different functional organization in parietal reach region and the dorsal premotor cortex. American Journal of Physiology-Heart and Circulatory Physiology doi:10.1152/jn.00621.2013

Chicharro, D., Kreuz, T., and Andrzejak, R. (2011). What can spike train distances tell us about the neural code? Journal of neuroscience methods 199, 146–165. doi:10.1016/j.jneumeth.2011.05.002

Cohen, M. and Kohn, A. (2011). Measuring and interpreting neuronal correlations. Nature neuroscience 14, 811. doi:10.1038/nn.2842

Cover, T. and Thomas, J. (2012). Elements of information theory (John Wiley & Sons). doi:10.1002/047174882X

Cowley, B., Kaufman, M., Butler, Z., Churchland, M., Ryu, S., Shenoy, K., et al. (2013). DataHigh: graphical user interface for visualizing and interacting with high-dimensional neural activity. Journal of neural engineering 10, 066012. doi:10.1088/1741-2560/10/6/066012

Cowley, B., Semedo, J., Zandvakili, A., Smith, M., Kohn, A., and Yu, B. (2017). Distance covariance analysis. In Artificial Intelligence and Statistics. 242–251

Cui, J., Xu, L., Bressler, S., Ding, M., and Liang, H. (2008). BSMART: a Matlab/C toolbox for analysis of multichannel neural time series. Neural Networks 21, 1094–1104. doi:10.1016/j.neunet.2008.05.007

Cunningham, J. and Byron, M. (2014). Dimensionality reduction for large-scale neural recordings. Nature neuroscience 17, 1500. doi:10.1038/nn.3776

Cutts, C. and Eglen, S. (2014). Detecting pairwise correlations in spike trains: an objective comparison of methods and application to the study of retinal waves. Journal of Neuroscience 34, 14288–14303. doi:10.1523/JNEUROSCI.2767-14.2014

Delorme, A., Mullen, T., Kothe, C., Acar, Z., Bigdely-Shamlo, N., Vankov, A., et al. (2011). EEGLAB, SIFT, NFT, BCILAB, and ERICA: new tools for advanced EEG processing. Computational intelligence and neuroscience 2011, 10. doi:10.1155/2011/130714

Deng, X., Eskandar, E., and Eden, U. (2013). A point process approach to identifying and tracking transitions in neural spiking dynamics in the subthalamic nucleus of Parkinson’s patients. Chaos: An Interdisciplinary Journal of Nonlinear Science 23, 046102. doi:10.1063/1.4818546

Egert, U., Knott, T., Schwarz, C., Nawrot, M., Brandt, A., Rotter, S., et al. (2002). MEA-Tools: an open source toolbox for the analysis of multi-electrode data with MATLAB. Journal of neuroscience methods 117, 33–42. doi:10.1016/S0165-0270(02)00045-6

Farge, M. (1992). Wavelet transforms and their applications to turbulence. Annual review of fluid mechanics 24, 395–458. doi:10.1146/annurev.fl.24.010192.002143

Fee, M., Mitra, P., and Kleinfeld, D. (1996). Automatic sorting of multiple unit neuronal signals in the presence of anisotropic and non-Gaussian variability. Journal of neuroscience methods 69, 175–188. doi:10.1016/S0165-0270(96)00050-7

Fries, P., Reynolds, J., Rorie, A., and Desimone, R. (2001). Modulation of oscillatory neuronal synchronization by selective visual attention. Science 291, 1560–1563. doi:10.1126/science.291.5508.1560

Garcia, S. and Fourcaud-Trocmé, N. (2009). OpenElectrophy: an electrophysiological data-and analysis-sharing framework. Frontiers in neuroinformatics 3, 14. doi:10.3389/neuro.11.014.2009

Garcia, S., Guarino, D., Jaillet, F., Jennings, T., Pröpper, R., Rautenberg, P., et al. (2014). Neo: an object model for handling electrophysiology data in multiple formats. Frontiers in neuroinformatics 8, 10. doi:10.3389/fninf.2014.00010

Geweke, J. (1982). Measurement of linear dependence and feedback between multiple time series. Journal of the American statistical association 77, 304–313. doi:10.2307/2287238

Glaser, J., Perich, M., Ramkumar, P., Miller, L., and Kording, K. (2018). Population coding of conditional probability distributions in dorsal premotor cortex. Nature communications 9, 1788. doi:10.1038/s41467-018-04062-6

Goldberg, D., Victor, J., Gardner, E., and Gardner, D. (2009). Spike train analysis toolkit: enabling wider application of information-theoretic techniques to neurophysiology. Neuroinformatics 7, 165–178. doi:10.1007/s12021-009-9049-y

Granger, C. (1969). Investigating causal relations by econometric models and cross-spectral methods. Econometrica: Journal of the Econometric Society, 424–438 doi:10.2307/1912791

Granlund, J. (1949). Interference in frequency-modulation reception

Grün, S., Diesmann, M., and Aertsen, A. (2002). Unitary events in multiple single-neuron spiking activity: I. detection and significance. Neural Computation 14, 43–80. doi:10.1162/089976602753284455

Grün, S., Diesmann, M., Grammont, F., Riehle, A., and Aertsen, A. (1999). Detecting unitary events without discretization of time. Journal of neuroscience methods 94, 67–79. doi:10.1016/S0165-0270(99)00126-0

Hafner, C. and Herwartz, H. (2008). Testing for causality in variance using multivariate GARCH models. Annales d’Economie et de Statistique, 215–241 doi:10.2307/27715168

Hayden, B., Smith, D., and Platt, M. (2009). Electrophysiological correlates of default-mode processing in macaque posterior cingulate cortex. Proceedings of the National Academy of Sciences 106, 5948–5953. doi:10.1073/pnas.0812035106

Hazan, L., Zugaro, M., and Buzsäki, G. (2006). Klusters, NeuroScope, NDManager: a free software suite for neurophysiological data processing and visualization. Journal of neuroscience methods 155, 207–216. doi:10.1016/j.jneumeth.2006.01.017

Hill, D., Mehta, S., and Kleinfeld, D. (2011). Quality metrics to accompany spike sorting of extracellular signals. Journal of Neuroscience 31, 8699–8705. doi:10.1523/JNEUROSCI.0971-11.2011

Holt, G., Softky, W., Koch, C., and Douglas, R. (1996). Comparison of discharge variability in vitro and in vivo in cat visual cortex neurons. Journal of neurophysiology 75, 1806–1814. doi:10.1152/jn.1996.75.5.1806

Hurtado, J., Rubchinsky, L., and Sigvardt, K. (2004). Statistical method for detection of phase-locking episodes in neural oscillations. Journal of neurophysiology 91, 1883–1898. doi:10.1152/jn.00853.2003

Hurtado, J., Rubchinsky, L., Sigvardt, K., Wheelock, V., and Pappas, C. T. (2005). Temporal evolution of oscillations and synchrony in GPi/muscle pairs in Parkinson’s disease. Journal of neurophysiology 93, 1569–1584. doi:10.1152/jn.00829.2004

Ince, R. (2012). Open-source software for studying neural codes. arXiv preprint arXiv:1207.5933 doi: 10.1201/b14756-35

Ince, R., Mazzoni, A., Petersen, R., and Panzeri, S. (2010). Open source tools for the information theoretic analysis of neural data. Frontiers in neuroscience 3, 11. doi:10.3389/neuro.01.011.2010

Ince, R., Petersen, R., Swan, D., and Panzeri, S. (2009). Python for information theoretic analysis of neural data. Frontiers in Neuroinformatics 3, 4. doi:10.3389/neuro.11.004.2009

Jarvis, M. and Mitra, P. (2001). Sampling properties of the spectrum and coherency of sequences of action potentials. Neural Computation 13, 717–749. doi:10.1162/089976601300014312

Kaminski, M. and Blinowska, K. (1991). A new method of the description of the information flow in the brain structures. Biological cybernetics 65, 203–210. doi:10.1007/BF00198091

Kobak, D., Brendel, W., Constantinidis, C., Feierstein, C., Kepecs, A., Mainen, Z., et al. (2016). Demixed principal component analysis of neural population data. Elife 5, e10989. doi:10.7554/eLife.10989

Kramer, M. and Eden, U. (2016). Case Studies in Neural Data Analysis: A Guide for the Practicing Neuroscientist (MIT Press)

Kreuz, T., Chicharro, D., Houghton, C., Andrzejak, R., and Mormann, F. (2012). Monitoring spike train synchrony. Journal of neurophysiology 109, 1457–1472. doi:10.1152/jn.00873.2012

Kreuz, T., Haas, J., Morelli, A., Abarbanel, H., and Politi, A. (2007). Measuring spike train synchrony. Journal of neuroscience methods 165, 151–161. doi:10.1016/j.jneumeth.2007.05.031

Kreuz, T., Mulansky, M., and Bozanic, N. (2015). SPIKY: A graphical user interface for monitoring spike train synchrony. Journal of neurophysiology 113, 3432–3445. doi:10.1152/jn.00848.2014

Lachaux, J.-P., Rodriguez, E., Martinerie, J., and Varela, F. (1999). Measuring phase synchrony in brain signals. Human brain mapping 8, 194–208. doi:10.1002/(SICI)1097-0193(1999)8:4〈194::AID-HBM4〉3.0.CO;2-C

Lakshmanan, K., Sadtler, P., Tyler-Kabara, E., Batista, A., and Yu, B. (2015). Extracting low-dimensional latent structure from time series in the presence of delays. Neural computation 27, 1825–1856. doi: 10.1162/NECO_a_00759

Lawlor, P., Perich, M., Miller, L., and Kording, K. (2018). Linear-nonlinear-time-warp-poisson models of neural activity. Journal of computational neuroscience 45, 173–191. doi:10.1007/s10827-018-0696-6

Le Van Quyen, M., Foucher, J., Lachaux, J.-P., Rodriguez, E., Lutz, A., Martinerie, J., et al. (2001). Comparison of Hilbert transform and wavelet methods for the analysis of neuronal synchrony. Journal of neuroscience methods 111, 83–98. doi:10.1016/S0165-0270(01)00372-7

Lidierth, M. (2009). sigTOOL: a MATLAB-based environment for sharing laboratory-developed software to analyze biological signals. Journal of neuroscience methods 178, 188–196. doi:10.1016/j.jneumeth.2008.11.0040

Lindner, M., Vicente, R., Priesemann, V., and Wibral, M. (2011). TRENTOOL: A Matlab open source toolbox to analyse information flow in time series data with transfer entropy. BMC neuroscience 12, 119. doi:10.1186/1471-2202-12-119

Liu, X.-Q., Wu, X., and Liu, C. (2011). SPKtool: An open source toolbox for electrophysiological data processing. In Biomedical Engineering and Informatics (BMEI), 2011 4th International Conference on (IEEE), vol. 2, 854–857. doi:10.1109/BMEI.2011.6098451

Lizier, J. (2014). JIDT: An information-theoretic toolkit for studying the dynamics of complex systems. Frontiers in Robotics and AI 1, 11. doi:10.3389/frobt.2014.00011

Loader, C. (2006). Local regression and likelihood (Springer Science & Business Media). doi:10.1007/b98858

[Dataset] Lowet, E., Roberts, M., Hadjipapas, A., Peter, A., van der Eerden, J., and De Weerd, P. (2015). Data from: Input-dependent frequency modulation of cortical gamma oscillations shapes spatial synchronization and enables phase coding. https://datadryad.org/resource/doi:10.5061/dryad.p631f. doi:https://doi.org/10.5061/dryad.p631f

Mackevicius, E., Bahle, A., Williams, A., Gu, S., Denisenko, N., Goldman, M., et al. (2019). Unsupervised discovery of temporal sequences in high-dimensional datasets, with applications to neuroscience. eLife 8, e38471. doi:10.7554/eLife.38471

Magri, C., Whittingstall, K., Singh, V., Logothetis, N., and Panzeri, S. (2009). A toolbox for the fast information analysis of multiple-site LFP, EEG and spike train recordings. BMC neuroscience 10, 81. doi:10.1186/1471-2202-10-81

Mahmud, M., Bertoldo, A., Girardi, S., Maschietto, M., and Vassanelli, S. (2012). SigMate: a Matlab-based automated tool for extracellular neuronal signal processing and analysis. Journal of neuroscience methods 207, 97–112. doi:10.1016/j.jneumeth.2012.03.009

Mahmud, M. and Vassanelli, S. (2016). Processing and analysis of multichannel extracellular neuronal signals: State-of-the-art and challenges. Frontiers in neuroscience 10, 248. doi:10.3389/fnins.2016.00248

Makino, H., Ren, C., Liu, H., Kim, A., Kondapaneni, N., Liu, X., et al. (2017). Transformation of cortex-wide emergent properties during motor learning. Neuron 94, 880–890. doi:10.1016/j.neuron.2017.04.015

Maris, E., Schoffelen, J.-M., and Fries, P. (2007). Nonparametric statistical testing of coherence differences. Journal of neuroscience methods 163, 161–175. doi:10.1016/j.jneumeth.2007.02.011

McFarland, J., Cui, Y., and Butts, D. (2013). Inferring nonlinear neuronal computation based on physiologically plausible inputs. PLoS computational biology 9, e1003143. doi:10.1371/journal.pcbi.1003143

Meier, R., Egert, U., Aertsen, A., and Nawrot, M. (2008). FIND – a unified framework for neural data analysis. Neural Networks 21, 1085–1093. doi:10.1016/j.neunet.2008.06.019

Mewett, D., Reynolds, K., and Nazeran, H. (2004). Reducing power line interference in digitised electromyogram recordings by spectrum interpolation. Medical and Biological Engineering and Computing 42, 524–531. doi:10.1007/BF02350994

Mitra, P. (2007). Observed brain dynamics (Oxford University Press)

Mitra, P. and Pesaran, B. (1999). Analysis of dynamic brain imaging data. Biophysical journal 76, 691–708. doi:10.1016/S0006-3495(99)77236-X

Montalto, A., Faes, L., and Marinazzo, D. (2014). MuTE: a MATLAB toolbox to compare established and novel estimators of the multivariate transfer entropy. PloS one 9, e109462. doi:10.1371/journal.pone.0109462

Moore, D., Valentini, G., Walker, S., and Levin, M. (2017). Inform: A toolkit for information-theoretic analysis of complex systems. In Computational Intelligence (SSCI), 2017 IEEE Symposium Series on (IEEE), 1–8. doi:10.1109/SSCI.2017.8285197

Morel, P. (2018). Gramm: grammar of graphics plotting for Matlab. The Journal of Open Source Software 3. doi:10.21105/joss.00568

Mulansky, M., Bozanic, N., Sburlea, A., and Kreuz, T. (2015). A guide to time-resolved and parameter-free measures of spike train synchrony. In Event-based Control, Communication, and Signal Processing (EBCCSP), 2015 International Conference on (IEEE), 1–8. doi:10.1186/1471-2202-16-S1-P133

Mullen, T. (2014). The dynamic brain: Modeling neural dynamics and interactions from human electrophysiological recordings (University of California, San Diego)

Niso, G., Bruña, R., Pereda, E., Gutiérrez, R., Bajo, R., Maestú, F., et al. (2013). Hermes: towards an integrated toolbox to characterize functional and effective brain connectivity. Neuroinformatics 11, 405–434. doi:10.1007/s12021-013-9186-1

Nolte, G., Bai, O., Wheaton, L., Mari, Z., Vorbach, S., and Hallett, M. (2004). Identifying true brain interaction from EEG data using the imaginary part of coherency. Clinical neurophysiology 115, 2292–2307. doi:10.1016/j.clinph.2004.04.029

Nolte, G., Ziehe, A., Nikulin, V., Schlögl, A., Krämer, N., Brismar, T., et al. (2008). Robustly estimating the flow direction of information in complex physical systems. Physical review letters 100, 234101. doi:10.1103/PhysRevLett.100.234101

Oostenveld, R., Fries, P., Maris, E., and Schoffelen, J.-M. (2011). FieldTrip: open source software for advanced analysis of MEG, EEG, and invasive electrophysiological data. Computational intelligence and neuroscience 2011, 1. doi:10.1155/2011/156869

Özkurt, T. and Schnitzler, A. (2011). A critical note on the definition of phase–amplitude cross-frequency coupling. Journal of Neuroscience methods 201, 438–443. doi:10.1016/j.jneumeth.2011.08.014

Pachitariu, M., Steinmetz, N., Kadir, S., Carandini, M., and Harris, K. (2016). Fast and accurate spike sorting of high-channel count probes with KiloSort. In Advances in Neural Information Processing Systems. 4448–4456

Paninski, L., Pillow, J., and Lewi, J. (2007). Statistical models for neural encoding, decoding, and optimal stimulus design. Progress in brain research 165, 493–507. doi:10.1016/S0079-6123(06)65031-0

Parikh, H. (2009). On Improving the Effectiveness of Control Signals from Chronic Microelectrodes for Cortical Neuroprostheses. Ph.D. thesis

Park, I., Meister, M., Huk, A., and Pillow, J. (2014). Encoding and decoding in parietal cortex during sensorimotor decision-making. Nature neuroscience 17, 1395. doi:10.1038/nn.3800

Park, I. and Pillow, J. (2011). Bayesian spike-triggered covariance analysis. In Advances in neural information processing systems. 1692–1700

Pastore, V., Poli, D., Godjoski, A., Martinoia, S., and Massobrio, P. (2016). ToolConnect: a functional connectivity toolbox for in vitro networks. Frontiers in neuroinformatics 10, 13. doi:10.3389/fninf.2016.00013

Percival, D. and Walden, A. (1993). Spectral analysis for physical applications (Cambridge University Press). doi:10.1017/CBO9780511622762

[Dataset] Perich, M., Lawlor, P., Miller, L., and Kording, K. (2018). Extracellular neural recordings from macaque primary and dorsal premotor motor cortex during a sequential reaching task. http://crcns.org/data-sets/motor-cortex/pmd-1. doi:10.6080/K0FT8J72

Pesaran, B. (2008). Spectral analysis for neural signals. In Short course III, presented at 2008 Society for Neuroscience Annual Meeting (Mitra P.P., ed.)

Pettersen, K., Devor, A., Ulbert, I., Dale, A., and Einevoll, G. (2006). Current-source density estimation based on inversion of electrostatic forward solution: effects of finite extent of neuronal activity and conductivity discontinuities. Journal of neuroscience methods 154, 116–133. doi:10.1016/j.jneumeth.2005.12.005

Pillow, J., Shlens, J., Paninski, L., Sher, A., Litke, A., Chichilnisky, E., et al. (2008). Spatio-temporal correlations and visual signalling in a complete neuronal population. Nature 454, 995. doi:10.1038/nature07140

Pouzat, C. and Chaffiol, A. (2009). Automatic spike train analysis and report generation. an implementation with r, r2html and star. Journal of neuroscience methods 181, 119–144. doi:10.1016/j.jneumeth.2009.01.037

Prechtl, J., Cohen, L., Pesaran, B., Mitra, P., and Kleinfeld, D. (1997). Visual stimuli induce waves of electrical activity in turtle cortex. Proceedings of the National Academy of Sciences 94, 7621–7626

Prieto, G., Parker, R., Thomson, D., Vernon, F., and Graham, R. (2007). Reducing the bias of multitaper spectrum estimates. Geophysical Journal International 171, 1269–1281

Pröpper, R. and Obermayer, K. (2013). Spyke Viewer: a flexible and extensible platform for electrophysiological data analysis. Frontiers in neuroinformatics 7, 26. doi:10.3389/fninf.2013.00026

Quaglio, P., Rostami, V., Torre, E., and Grün, S. (2018). Methods for identification of spike patterns in massively parallel spike trains. Biological cybernetics 112, 57–80. doi:10.1007/s00422-018-0755-0

Quaglio, P., Yegenoglu, A., Torre, E., Endres, D., and Grün, S. (2017). Detection and evaluation of spatiotemporal spike patterns in massively parallel spike train data with SPADE. Frontiers in computational neuroscience 11, 41. doi:10.3389/fncom.2017.00041

Quiroga, R., Kreuz, T., and Grassberger, P. (2002). Event synchronization: a simple and fast method to measure synchronicity and time delay patterns. Physical review E 66, 041904. doi:10.1103/PhysRevE.66.041904

Quiroga, R., Nadasdy, Z., and Ben-Shaul, Y. (2004). Unsupervised spike detection and sorting with wavelets and superparamagnetic clustering. Neural computation 16, 1661–1687. doi:10.1162/089976604774201631

Rankine, L., Stevenson, N., Mesbah, M., and Boashash, B. (2005). A quantitative comparison of non-parametric time-frequency representations. In 2005 13th European Signal Processing Conference (IEEE), 1–4

Rivlin-Etzion, M., Ritov, Y., Heimer, G., Bergman, H., and Bar-Gad, I. (2006). Local shuffling of spike trains boosts the accuracy of spike train spectral analysis. Journal of neurophysiology 95, 3245–3256. doi:10.1152/jn.00055.2005

Rosenberg, J., Amjad, A., Breeze, P., Brillinger, D., and Halliday, D. (1989). The Fourier approach to the identification of functional coupling between neuronal spike trains. Progress in biophysics and molecular biology 53, 1–31. doi:10.1016/0079-6107(89)90004-7

Rosenberg, J., Halliday, D., Breeze, P., and Conway, B. (1998). Identification of patterns of neuronal connectivitypartial spectra, partial coherence, and neuronal interactions. Journal of neuroscience methods 83, 57–72. doi:10.1016/S0165-0270(98)00061-2

Russo, E. and Durstewitz, D. (2017). Cell assemblies at multiple time scales with arbitrary lag constellations. Elife 6, e19428. doi:10.7554/eLife.19428

Samiee, S. and Baillet, S. (2017). Time-resolved phase-amplitude coupling in neural oscillations. NeuroImage 159, 270–279. doi:10.1016/j.neuroimage.2017.07.051

Schrader, S., Grun, S., Diesmann, M., and Gerstein, G. (2008). Detecting synfire chain activity using massively parallel spike train recording. Journal of neurophysiology 100, 2165–2176. doi:10.1152/jn.01245.2007

Scott, D. (1979). On optimal and data-based histograms. Biometrika 66, 605–610. doi:10.1093/biomet/66.3.605

Shimazaki, H. and Shinomoto, S. (2010). Kernel bandwidth optimization in spike rate estimation. Journal of computational neuroscience 29, 171–182. doi:10.1007/s10827-009-0180-4

Shinomoto, S., Miura, K., and Koyama, S. (2005). A measure of local variation of inter-spike intervals. Biosystems 79, 67–72. doi:10.1016/j.biosystems.2004.09.023

Shinomoto, S., Shima, K., and Tanji, J. (2003). Differences in spiking patterns among cortical neurons. Neural Computation 15, 2823–2842. doi:10.1162/089976603322518759

Shlens, J. (2014). A tutorial on principal component analysis. arXiv preprint arXiv:1404.1100

Slepian, D. and Pollak, H. (1961). Prolate spheroidal wave functions, Fourier analysis and uncertaintyI. Bell System Technical Journal 40, 43–63. doi:10.1002/j.1538-7305.1964.tb01037.x

Stam, C., Nolte, G., and Daffertshofer, A. (2007). Phase lag index: assessment of functional connectivity from multi channel EEG and MEG with diminished bias from common sources. Human brain mapping 28, 1178–1193. doi:10.1002/hbm.20346

Staude, B., Rotter, S., and Grün, S. (2010). CuBIC: cumulant based inference of higher-order correlations in massively parallel spike trains. Journal of computational neuroscience 29, 327–350. doi:10.1007/s10827-009-0195-x

Stevenson, I. and Kording, K. (2011). How advances in neural recording affect data analysis. Nature neuroscience 14, 139. doi:10.1038/nn.2731

Stoica, P., Moses, R., et al. (2005). Spectral analysis of signals (Pearson Prentice Hall Upper Saddle River, NJ)

Tadel, F., Baillet, S., Mosher, J., Pantazis, D., and Leahy, R. (2011). Brainstorm: a user-friendly application for MEG/EEG analysis. Computational intelligence and neuroscience 2011, 8. doi:10.1155/2011/879716

Tallon-Baudry, C., Bertrand, O., Delpuech, C., and Pernier, J. (1997). Oscillatory *γ*-band (30-70 hz) activity induced by a visual search task in humans. Journal of Neuroscience 17, 722–734. doi:10.1523/JNEUROSCI.17-02-00722.1997

Thomson, D. (1982). Spectrum estimation and harmonic analysis. Proceedings of the IEEE 70, 1055–1096. doi:10.1109/PROC.1982.12433

Timme, N. and Lapish, C. (2018). A tutorial for information theory in neuroscience. eNeuro 5. doi: 10.1523/ENEURO.0052-18.2018

Torre, E., Canova, C., Denker, M., Gerstein, G., Helias, M., and Grün, S. (2016). ASSET: analysis of sequences of synchronous events in massively parallel spike trains. PLoS computational biology 12, e1004939. doi:10.1371/journal.pcbi.1004939

Torre, E., Picado-Muiño, D., Denker, M., Borgelt, C., and Grün, S. (2013). Statistical evaluation of synchronous spike patterns extracted by frequent item set mining. Frontiers in computational neuroscience 7, 132. doi:10.3389/fncom.2013.00132

Tort, A., Komorowski, R., Eichenbaum, H., and Kopell, N. (2010). Measuring phase-amplitude coupling between neuronal oscillations of different frequencies. Journal of neurophysiology 104, 1195–1210. doi:10.1152/jn.00106.2010

Truccolo, W., Eden, U., Fellows, M., Donoghue, J., and Brown, E. (2005). A point process framework for relating neural spiking activity to spiking history, neural ensemble, and extrinsic covariate effects. Journal of neurophysiology 93, 1074–1089. doi:10.1152/jn.00697.2004

van Rossum, M. (2001). A novel spike distance. Neural computation 13, 751–763. doi:10.1162/089976601300014321

van Vugt, M., Sederberg, P., and Kahana, M. (2007). Comparison of spectral analysis methods for characterizing brain oscillations. Journal of neuroscience methods 162, 49–63. doi:10.1016/j.jneumeth.2006.12.004

Victor, J. and Purpura, K. (1996). Nature and precision of temporal coding in visual cortex: a metric-space analysis. Journal of neurophysiology 76, 1310–1326. doi:10.1152/jn.1996.76.2.1310

Vinck, M., Battaglia, F., Womelsdorf, T., and Pennartz, C. (2012). Improved measures of phase-coupling between spikes and the local field potential. Journal of computational neuroscience 33, 53–75. doi: 10.1007/s10827-011-0374-4

Vinck, M., Oostenveld, R., Van Wingerden, M., Battaglia, F., and Pennartz, C. (2011). An improved index of phase-synchronization for electrophysiological data in the presence of volume-conduction, noise and sample-size bias. Neuroimage 55, 1548–1565. doi:10.1016/j.neuroimage.2011.01.055

Vinck, M., van Wingerden, M., Womelsdorf, T., Fries, P., and Pennartz, C. (2010). The pairwise phase consistency: a bias-free measure of rhythmic neuronal synchronization. Neuroimage 51, 112–122. doi:10.1016/j.neuroimage.2010.01.073

Voytek, B., Canolty, R., Shestyuk, A., Crone, N., Parvizi, J., and Knight, R. (2010). Shifts in gamma phase-amplitude coupling frequency from theta to alpha over posterior cortex during visual tasks. Frontiers in human neuroscience 4, 191. doi:10.3389/fnhum.2010.00191

Wagenaar, D., DeMarse, T., and Potter, S. (2005). MeaBench: A toolset for multi-electrode data acquisition and on-line analysis. In Neural Engineering, 2005. Conference Proceedings. 2nd International IEEE EMBS Conference on (IEEE), 518–521. doi:10.1109/CNE.2005.1419673

Williams, A., Kim, T., Wang, F., Vyas, S., Ryu, S., and Shenoy, K. e. a. (2018). Unsupervised discovery of demixed, low-dimensional neural dynamics across multiple timescales through tensor component analysis. Neuron doi:10.1016/j.neuron.2018.05.015

Wong, R., Meister, M., and Shatz, C. (1993). Transient period of correlated bursting activity during development of the mammalian retina. Neuron 11, 923–938. doi:10.1016/0896-6273(93)90122-8

Yegenoglu, A., Holstein, D., Phan, L., Denker, M., Davison, A., and Grün, S. (2017). Elephant–open-source tool for the analysis of electrophysiological data sets. In Bernstein Conference 2015: Abstract Book. W–05. doi:10.12751/nncn.bc2015.0126

Yu, B., Cunningham, J., Santhanam, G., Ryu, S., Shenoy, K., and Sahani, M. (2009). Gaussian-process factor analysis for low-dimensional single-trial analysis of neural population activity. In Advances in neural information processing systems. 1881–1888. doi:10.1152/jn.90941.2008

